# Liver Kinome Profiling Identifies PS1145 as a Potential Therapeutic molecule for amelioration of Systemic Inflammation in Alcohol-related Liver Disease

**DOI:** 10.1101/2025.07.02.662754

**Authors:** Manisha Yadav, Abhishak Gupta, Sanju Yadav, Vipul Sharma, Yash Magar, Deepanshu Deepanshu, Tahseen Khan, Neha Sharma, Kavita Yadav, Nupur Sharma, Vasundhra Bindal, Sushmita Pandey, Gaurav Tripathi, Rimsha Saif, Babu Mathew, Dinesh Mani Tripathi, Anupama Kumari, Shiv Kumar Sarin, Jaswinder Singh Maras

**Affiliations:** Department of Molecular and Cellular Medicine, Institute of Liver and Biliary Sciences, New Delhi; Centre of Comparative Medicine, Institute of Liver and Biliary Sciences, New Delhi; Amity University, Gurgaon; Artemis Hospital, Gurgaon; Department of Hepatology, Institute of Liver and Biliary Sciences, New Delhi; National Institute of Immunology, New Delhi

**Keywords:** Phosphoproteomics, Alcohol-related Liver Disease, Kinome signaling, inflammation, PS1145, PH787904, Prednisolone, pre-clinical ALD Rat model

## Abstract

**Background and aims:** Alcohol-related liver disease (ALD) has high mortality due to systemic inflammation. We analysed liver and monocyte kinome profiles in a chronic ethanol-fed pre-clinical rat model to identify therapeutic targets that could mitigate inflammation in ALD.

**Method:** Kinome profile was performed in liver and circulating monocytes at baseline, and after 8,12,16,20 and 24-weeks of 40% ethanol administration. Pathway-specific inhibitors, including PS1145 (IKK-phosphorylation inhibitor), PH-797804 (MAPK14 inhibitor), resveratrol and prednisolone were tested for their anti-inflammatory effects in ALD-rats, PBMC from patients and NIAAA mouse model.

**Results:** Kinome profiling identified 497 liver and 345 monocyte kinases in ALD rats (FDR<0.01). A time-dependent increase in MAPK14-associated kinases was observed in both tissues (FC>1.5, p<0.05). By 24 weeks, 172 liver and 48 monocyte kinases were significantly upregulated, particularly those linked to MyD88–TLR4, PI3K–Akt, TNF, TGFβ, and cellular senescence pathways. Key contributors included TGFβR1, ROS-generating kinases, and IL-1-driven MAPK14 and IKK phosphorylation. Targeting this axis, PS1145 (an IKK inhibitor) suppressed NFκB activation and inflammation in THP1 and HepG2 cells, as well as PBMCs from healthy and SAH patients, outperforming PH797804, resveratrol, and prednisolone (p<0.05, FC>1.5). PS1145 significantly reduced IL-6, TNFα, and NFκB, while increasing IL-10. In vivo, PS1145 treatment in the NIAAA mouse model markedly reduced hepatic steatosis, cellular stress and inflammatory pathways (cytokine signalling, IL-36 signalling and others) more effectively than standard therapies, Notably, it also downregulated IL36R, impairing TLR receptor dimerization, suggesting a dual mechanism of action (p<0.05) highlighting its therapeutic potential in ameliorating systemic inflammation in ALD.

**Conclusion:** Our study highlights the key role of MAPK14 and MYD88–TLR4 pathway kinases in driving systemic inflammation in SAH. The IKK inhibitor PS1145, by blocking both IKK phosphorylation and TLR dimerization, effectively suppresses inflammatory signalling and improves liver pathology, positioning it as a promising targeted therapy for ALD.

## INTRODUCTION

Alcohol-related liver disease (ALD) is a major global health burden, driven by the chronic inflammation and immune dysregulation caused by excessive alcohol consumption [1]. Persistent alcohol exposure results in hepatocellular injury triggering an inflammatory response by recruiting and activating immune cells [2] which releases pro-inflammatory cytokines, exacerbating liver damage and creating a vicious cycle of inflammation and tissue injury [2]. Over time, this chronic inflammation promotes fibrosis, impairs liver function, and can progress to cirrhosis or hepatocellular carcinoma [3]. Thus, controlling systemic inflammation may help mitigate ALD progression and improve outcomes.

Treatment options for ALD are limited, with abstinence as the key strategy. However, severe alcohol-related hepatitis (SAH) often necessitates medical intervention [4]. Corticosteroids are the primary pharmacological treatment for SAH; however, many patients fail to respond, highlighting the urgent need for alternative therapies [5]. The lack of targeted treatments for ALD emphasizes the importance of understanding the molecular mechanisms underlying disease progression [6].

Kinases, as pivotal regulators of cellular signalling, play a central role in mediating inflammatory and stress responses in ALD [7]. Among these, the MAPK-pathway is a key signalling cascade activated by alcohol-induced stress [8]. Kinases such as MAPK-14 (p38) and NF-κB are crucial mediators that regulate the transcription of pro-inflammatory cytokines, perpetuating hepatic inflammation [9]. Although MAPK kinase inhibitors have been investigated in preclinical and clinical settings, studies have shown limited or no therapeutic efficacy in the treatment of alcoholic liver disease [8]. Investigating the dynamics of kinase signalling in ALD can provide valuable insights into temporal changes in these pathways and identify potential therapeutic targets.

Advances in mass spectrometry-based phosphoproteomics enable high-throughput analysis of phosphorylation events regulating protein function. This technology offers a comprehensive approach to studying the kinome and its alterations in disease states [10]. By leveraging this approach, it becomes possible to pinpoint dysregulated kinases and pathways that drive ALD progression [11].

The current study profiles kinome dynamics in the liver and circulating monocytes of an ALD-rat model across multiple time points during disease progression. Temporal analysis revealed key kinases, including IκB and MAPK14, whose dysregulation was closely associated with inflammatory responses. Specifically, IKK phosphorylates IκBα, leading to its degradation and subsequent translocation of NF-κB to the nucleus, where it promotes the transcription of inflammatory genes [12]. Similarly, MAPK14 is activated by stress stimuli and inflammatory cytokines, further driving the expression of pro-inflammatory mediators [13]. Importantly, the application of phosphoproteomics in this context enables comprehensive, unbiased mapping of phosphorylation events, providing critical insights into the dynamic regulation of signalling pathways during ALD progression.

To assess therapeutic potential, we evaluated the efficacy of PS1145 (an IKK inhibitor) and PH797804 (a MAPK14 inhibitor) *in-vitro* using THP1, HepG2, and PBMCs (from healthy donors and SAH patients), as well as *in-vivo* using the NIAAA model. Both agents were compared against resveratrol and prednisolone. PS1145 demonstrated robust suppression of inflammatory signalling by inhibiting IKK phosphorylation and reducing NF-κB activation, supporting its potential as a targeted therapy for systemic inflammation in SAH.

Importantly, to our knowledge, no previous studies have directly compared the efficacy of PS1145 and PH797804 in models of alcoholic liver disease. Our study therefore provides novel insight by directly evaluating and contrasting these two targeted approaches in ALD, supporting further investigation of IKK inhibition as a therapeutic strategy in this disease.

## MATERIAL AND METHODS

### Animal experimentation

All animal procedures were approved by the Institutional Animal Ethics Committee (IAEC), with the approval code IAEC/ILBS/007 for the ethanol administration to rats and for developing NIAAA mice model under Approval No: IAEC/ILBS/23/1. Human samples enrolled in the study were approved by Institutional Review Board-IEC/2022/93/MA08.

### ALD rat model development

ALD rat model was developed by feeding 40% ethanol diet for a period of 6 months as detailed in supplementary methods.

Hematoxylin and eosin (HE), oil red stain (OR), and Sirius Red (SR) staining were performed as per the standard protocol to analyze immune infiltration, lipid accumulation, and fibrosis, further detailed in supplementary methods. Plasma was collected to analyse liver function tests and lipid profiles, while PBMCs were isolated using a density gradient protocol. Liver tissue samples and PBMCs were preserved in liquid nitrogen for future analysis. Monocytes were isolated from blood as detailed in supplementary methods.

### Kinome Identification

To identify and quantify global kinases in rat liver and in circulating monocytes, we performed phosphoproteomics using high resolution mass spectrometry (Q Exactive Orbitrap, Thermo) technique. Raw data files were read in proteome discoverer software using Mascot and SequestHT. ptmRS was incorporated in the workflow to identify the phosphorylation status and position in all samples. All samples were pooled group wise and measured in triplicate reducing the analytical variance. Coral (https://phanstiel-lab.med.unc.edu/CORAL/) was used to identify and map the kinases in our data set. Major Kinome figures were generated using Coral map.

### Proteomics sample preparation

Liver tissue lysates (50 µg protein) were reduced with 10 mM DTT (60°C, 1 hr), alkylated with 10 mM IAA (RT, dark), and digested with trypsin (24 hr, 37°C). Peptides were desalted using C18 spin columns, lyophilized, and reconstituted in 0.1% formic acid. Nano-electrospray ionization MS/MS was performed using a Q-Exactive Plus instrument. Peptides were enriched on a trap column, separated on an analytical column, and eluted over a 120-min gradient. Mass spectra were acquired at 70,000 resolution, and protein identification was performed using Mascot, with FDR set at 0.01 and significance at p<0.05 as detailed in supplementary methods [14].

### Phosphoproteomics sample preparation

Compared to proteomics large amount protein samples (1mg) is required in phosphoproteomics. Therefore, pooling strategy was adopted to reach the required amount of protein for the phosphoproteomics. 1 mg protein sample was prepared same as proteomics sample. Post elution from C18 columns samples were lyophilized and were stored. 3 samples were pooled together and were reconstituted in the binding buffer (provided in the kit) for phosphoproteomics. TiO2 (Thermo, A32993) and Fe-NTA phospho-peptide enrichment kit (Thermo, A32992) were used for phosphoprotein enrichment. Samples in binding buffer were passed from TiO2/Fe-NTA column by following standard protocol mentioned in the kit manual. Eluted samples were again lyophilized and were reconstituted in 0.1% formic acid [15].

#### Validation experiments

For the validation experiments THP1 cells, HepG2 cells, SAH patients and healthy donor PBMC were used as *in-vitro* models system and NIAAA binge model as *in-vivo* approach. Maintenance protocol of THP1 and HepG2 culture system is provided in detailed in supplementary section. MTT assay was performed to evaluate the toxicity or tolerated dose of PS1145 (Cat. No.: HY-18008, Medchem), PH787904 (Cat. No.: HY-10403), and Resveratrol [16] in the *in-vitro* system (detailed in supplementary figure 5). Anti-inflammatory effect of 70nM PS1145, 30nM PH797804, and 25uM Resveratrol on the culture system was evaluated using qRT-PCR and proteomics as discussed in supplementary method. The results were further validated in PBMCs of SAH patients and healthy donor. Histological changes and molecular changes induced in liver by these compounds were finally assessed in NIAAA binge mouse model.

### Evaluation of Anti-Inflammatory Effects of PS1145, PH797804, Resveratrol and Prednisolone

THP-1-derived macrophages and HepG2 cells were activated with 150 mM ethanol, 100 ng/mL LPS, a combination of both, or 10% SAH patient plasma for 24 hours. Untreated controls were included. Cells were then treated with PS1145 (70 nM), PH797804 (30 nM), resveratrol (25 µM), or prednisolone (10 µM) for 24 hours. Inflammatory responses were assessed using qPCR and proteomics analysis.

### Evaluation of Anti-Inflammatory Effects of PS1145, PH797804, Resveratrol and Prednisolone in primary PBMC

Healthy PBMC (n=6) and SAH PBMCs (n=12) were procured from National Liver disease biobank, New Delhi. Cells were washed with PBS and then equal number of cells were seed in each well and incubated overnight in RPMI media. Healthy PBMC were tweaked with 100ng/ml LPS for 4 hours and were then treated with 70nM PS1145, 30nM PH797804, 25uM Resveratrol and 10uM Prednisolone. Samples were then subjected to qPCR analysis.

#### SAH patient selection criteria

Patients diagnosed with severe alcohol-related hepatitis (SAH) were included based on: (1) history of chronic alcohol use, (2) recent onset of jaundice, (3) Maddrey’s Discriminant Function (DF) >32, and (4) serum bilirubin >3 mg/dL. Patients with active infections, hepatocellular carcinoma, or other chronic liver diseases were excluded.

#### Mouse model of chronic and binge ethanol feeding (the NIAAA model)

Male C57BL/6 mice were subjected to the NIAAA model of chronic and binge ethanol feeding. Mice were fed a Lieber-DeCarli liquid diet containing 5% (v/v) ethanol ad libitum for 10 days to induce chronic ethanol exposure. From days 8 to 11, mice received a single daily dose of either PS1145 (50 mg/kg, intraperitoneal), PH797804 (50 mg/kg, intraperitoneal), or prednisolone (5 mg/kg, oral) administered. On day 11, ethanol-fed mice were given a single binge dose of ethanol (5 g/kg body weight) by oral gavage, while control mice received an isocaloric maltose dextrin gavage. Mice were euthanized 9 hours after the binge dose for sample collection. This protocol allowed evaluation of the therapeutic efficacy of the compounds during the progression of alcoholic liver injury [17].

### Pathway activity

Pathway activity of pathways enriched in proteomics dataset was calculated by taking average of proteins enriched in particular pathway.

### Statistical analysis

For proteomics, initial data underwent analysis using Proteome Discoverer version 2.3 (Thermo Fisher Scientific, USA). Statistical analyses were conducted using MetaboAnalyst 6.0 and GraphPad Prism-6. For comparisons between the two groups, unpaired (2-tailed) Student t-tests and Mann-Whitney U tests were employed. ANOVA and Kruskal-Wallis tests were used for comparisons among multiple groups. Correlations were assessed using Spearman correlation analysis, with coefficients (R²) greater than 0.5 considered significant. Features with p< 0.05 are considered significant.

## RESULTS

### ALD rat model development

An ALD-rat model was developed using Long Evans rats fed with 40% ethanol LDC diet for 24 weeks. Rats were sacrificed at 8, 12, 16, 20, and 24 weeks for kinome profiling in liver tissue and monocytes. Model validation was performed using histology, biochemical analysis, and mass spectrometry of alcohol metabolism proteins, ALD markers, interleukins, and mitochondrial stress proteins (Figure-1A).

**Figure 1:**
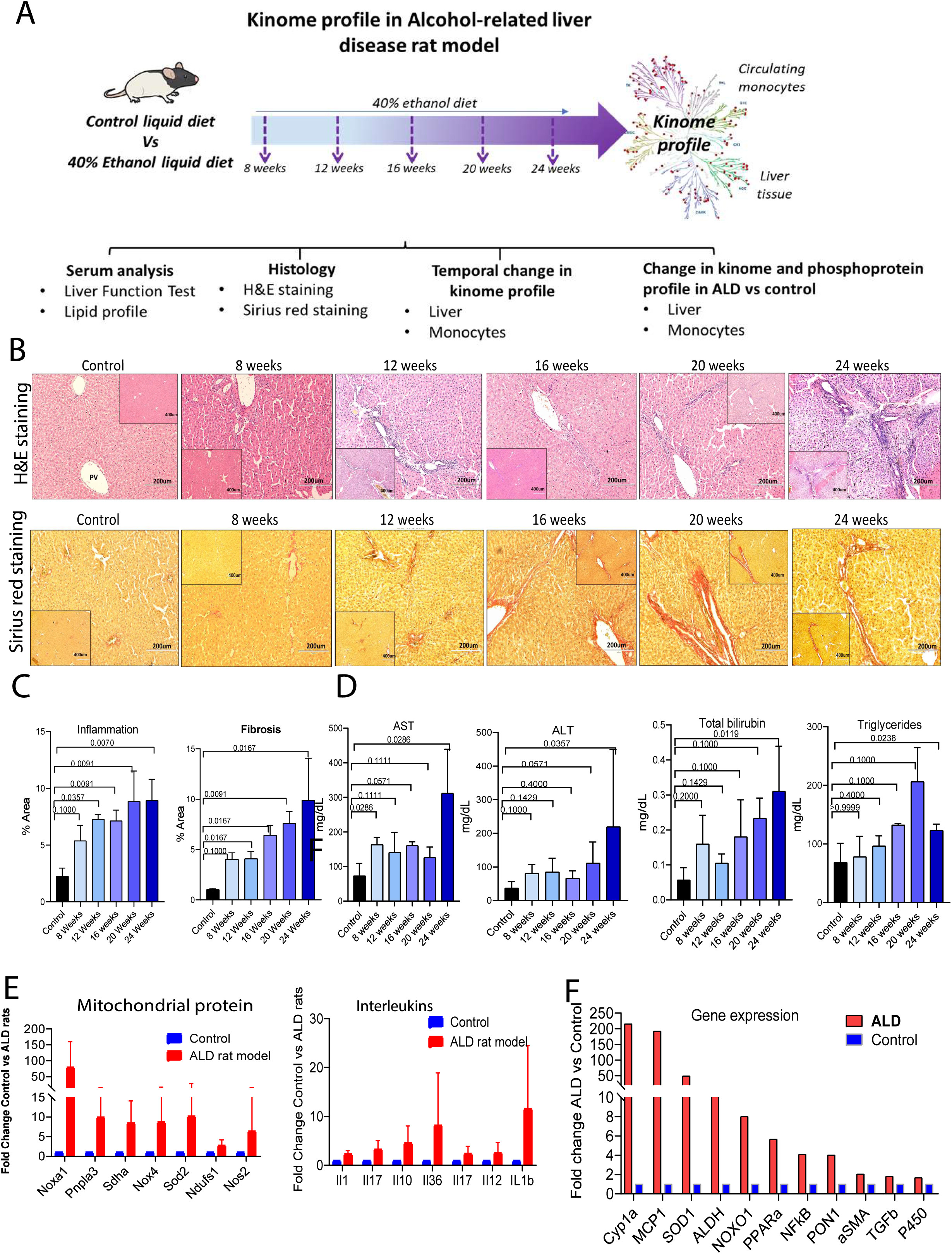
ALD rat model development. **A**: Study design. Rats were fed with 40% ethanol for 24 weeks. Serum was analysed for LFT and lipid profile. Histopathological analysis was performed in liver. Liver tissue and circulating cells at all time points were subjected to kinome profile. Identified were validated in *in-vitro* system. **B**: H&E and Sirius red staining of liver tissue of control, ALD rats at 8 weeks, 12 weeks, 16 weeks, 20 weeks and 24 weeks showing neutrophil infiltration and collagen deposition respectively. **C**: Scoring of H&E and Sirius red staining. % area of inflammation and fibrosis increased temporally over the time. **D**: Biochemical parameters showing AST, ALT, total bilirubin, and triglycerides. **E**: Bar plot showing increase in mitochondrial stress associated proteins and interleukins in ALD vs control **F**: Bar plot showing increase in inflammatory, alcohol metabolism genes and others in ALD vs control.

Histopathological analysis confirmed inflammation as a key feature of progressive alcoholic liver injury. H&E staining showed increasing immune cell infiltration, while Sirius Red staining revealed fibrosis progression, peaking at 24 weeks (Figure-1B). Quantitative scoring confirmed significant inflammation and fibrosis (Figure-1C).

Biochemical analysis showed elevated AST by week 8, ALT by week 20, and bilirubin by week 24, with triglycerides rising until week 20 before declining (Figure-1D). Proteomic analysis at 24 weeks highlighted a significant increase in mitochondrial stress proteins (Noxa1, Pnpla3, Sdha, Nox4, Sod2) (Figure-1E) and upregulation of alcohol metabolism and ALD-related proteins (Cyp1A, Aldh, Pon1, NF-κb, Tgf-β) (Figure-1F). These findings confirm the successful establishment of the ALD-rat model, providing a robust platform for studying kinome alterations associated with ALD.

### Comprehensive Kinome Profiling and Subcellular Localization in Rodent Liver and Monocytes

Liver tissue and circulating monocytes were subjected to a comprehensive global phosphoproteomic analysis to identify and quantify the kinases expressed in these samples. Ultra high-performance liquid chromatography coupled with mass spectrometry (UHPLC-MS) identified a total of 19,313 phospoproteins in the liver and 6,063 phosphoproteins in the monocytes. To date, 536 kinases have been documented (Human Kinome Database, Coral Kinome, Cell Signaling Technology, and KinMap) of which 497 kinases were identified in the liver and 345 kinases in monocytes (Figure-2A). Class-wise kinase distribution in liver and monocytes is shown in Figure-2B and phylogenetically mapped in Figure-2C.

**Figure 2.**
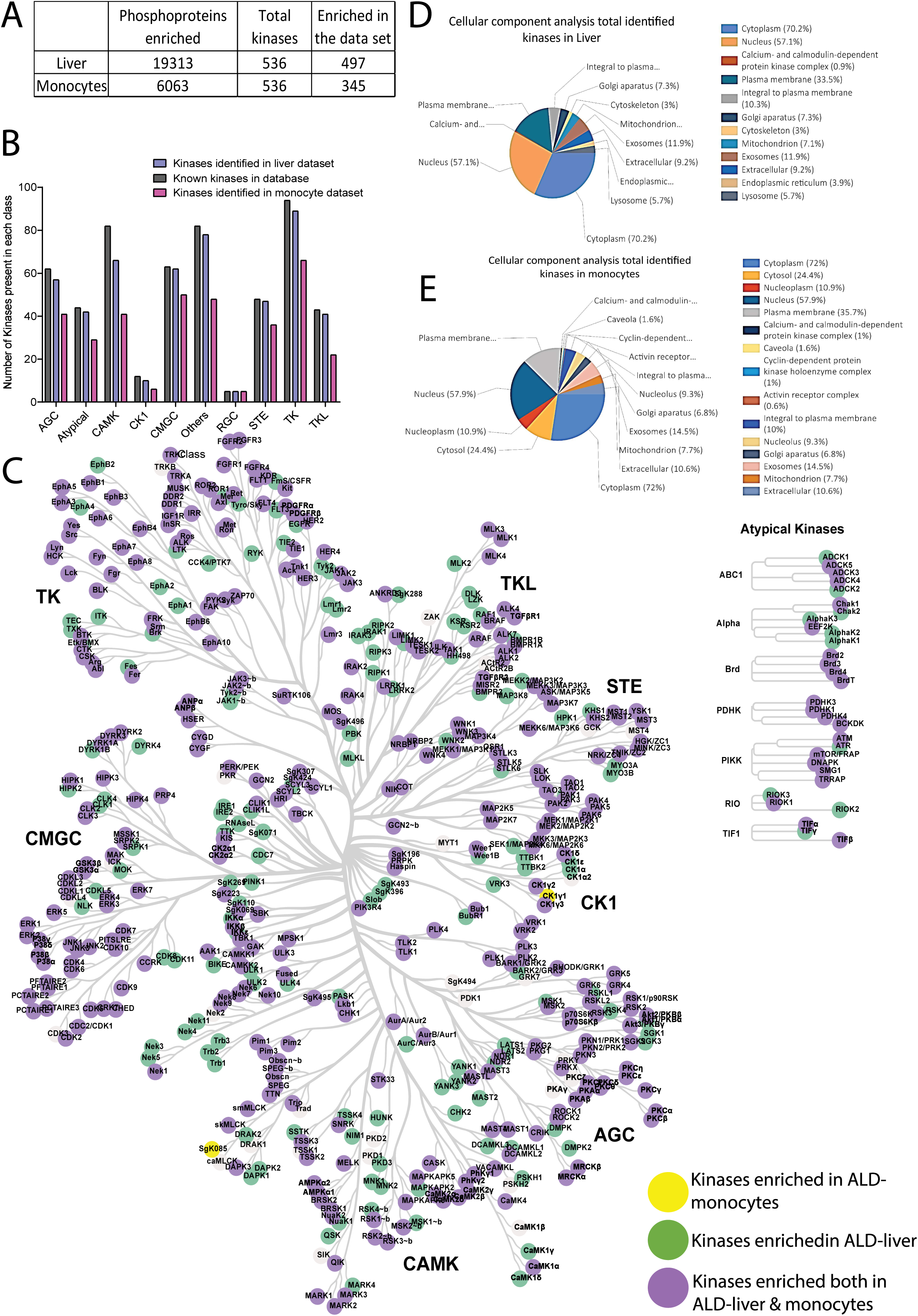
Comprehensive Kinome Profiling and Subcellular Localization in Rodent Liver and Monocytes. **A:** Enriched phosphoproteins, and kinases in liver and circulating monocyte datasets **B:** Class wise distribution of identified kinases in liver and circulating monocytes. **C:** Cellular component analysis of identified kinases in liver dataset. **D:** Cellular component analysis of identified kinases in circulating monocyte dataset. **E:** Kinome map representing identified kinases in liver and monocytes. Kinases present both in liver and circulating monocytes are highlighted in purple color, kinases present exclusively in liver and monocytes are highlighted in green and yellow colour respectively.

Subcellular analysis revealed kinase localization in liver and monocytes. In the liver, kinases were mainly in the cytoplasm-306, nucleus-249, plasma membrane-146, Golgi-32, ER-17, and mitochondria-31 (Figure-2D). Monocyte distribution included the cytoplasm-224, nucleus-180, plasma membrane-111, Golgi-21, ER-10, and mitochondria-24 and others in additional compartments, (Figure-2E, supplementary figure-1 and 2).

Together, these findings provide a detailed kinome profile and subcellular localization map for rodent liver and monocytes, serving as a valuable resource for understanding kinase signalling and pathways in ALD.

### Kinome profile of ALD-rat liver

Kinases are central regulators of signal transduction pathways and play pivotal roles in controlling critical cellular processes such as inflammation, apoptosis, and fibrosis [18]. To elucidate the alterations in kinase expression associated with ALD progression, a global kinome profile was performed in ALD-rat liver tissues. A total of 497 kinases, providing a comprehensive overview of kinase dysregulation in ALD-rats were identified and quantitated. Principal Component Analysis (PCA) revealed clear differences in the liver kinome profile of ALD-rats and controls (Figure-3A). Among the 497 kinases, 169 were significantly up- and 85 were significantly downregulated in ALD-rat liver compared to healthy (Figure-3B, Supplementary Table-1). The class-wise changes in kinase expression are shown in Figure-3C.

**Figure 3:**
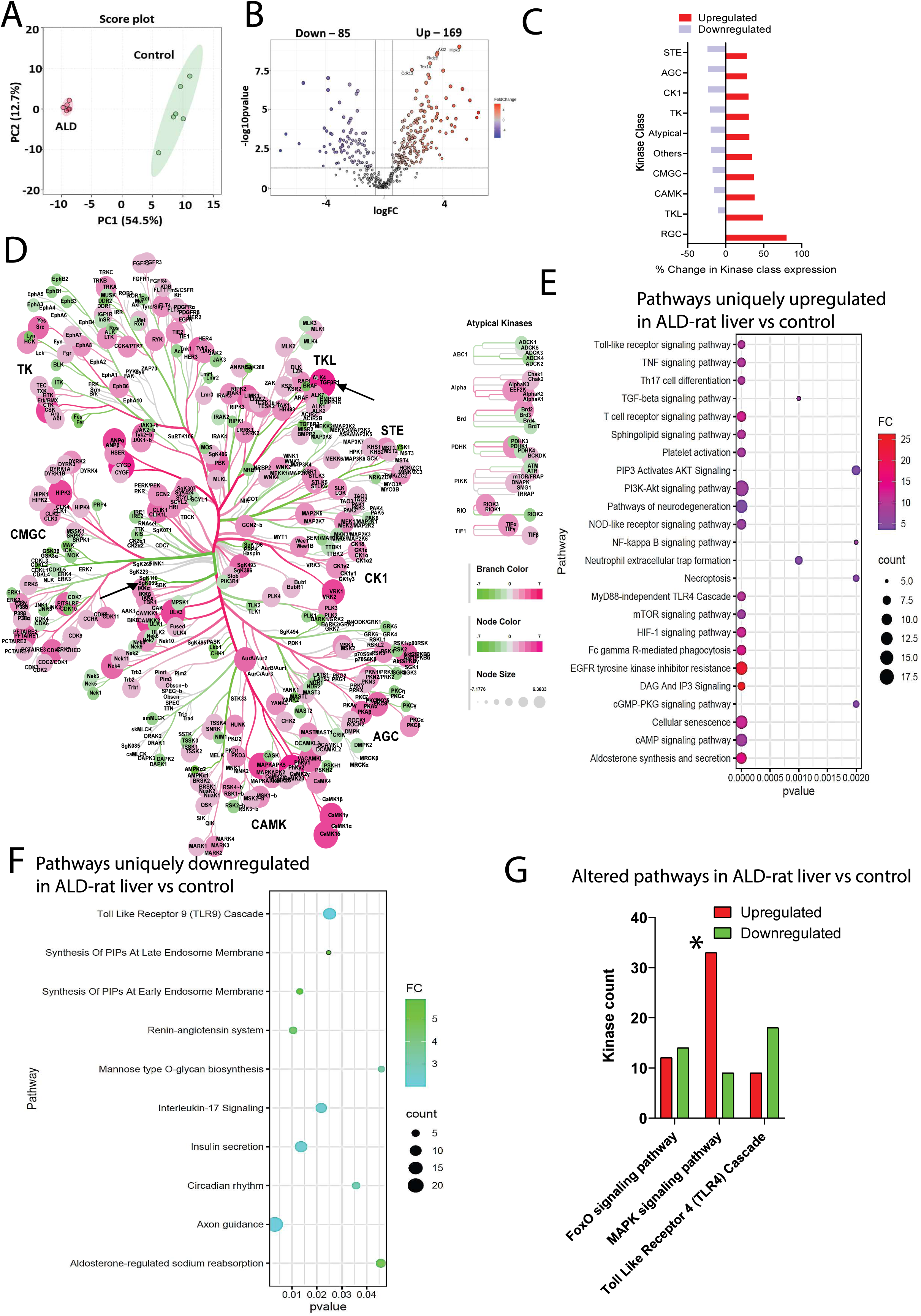
Kinome profile of ALD-rat liver. **A:** Principle component analysis showing segregation in liver kinome profile of ALD rats and control rats at 24 weeks. **B:** Volcano plot showing differential kinases in ALD rat liver compared to healthy rats. 169 kinases were significantly upregulated and 85 kinases were significantly downregulated in ALD rats. **C**: Bar plot showing percentage change in kinase class expression in ALD liver compared to healthy control liver. **D:** Kinome map representing differential kinases in ALD liver. Pink= high expression, green: low expression. **E:** Dot plot showing pathways significantly upregulated in ALD liver compared to control. **F:** Dot plot showing pathways significantly downregulated in ALD liver compared to control. **G:** bar plot showing altered pathways in ALD liver (involving both upregulated and downregulated kinases). 33 kinases associated with MAPK signaling were upregulated.

A kinome tree map illustrated the differentially expressed kinases in ALD-rat liver tissue. TGFBR1 kinase was the most up-regulated kinases in the liver (83.4-fold increase), suggesting fibrosis progression in ALD-rats (p<0.05, Figure-3D). Other top differentially upregulated kinases in ALD-rat liver were GUCY2D, PHKG2, CAMK1D, ULK3 and others (p<0.05, FC>1.5). Conversely, SBK2 (Sgk069), with essential role in regulating hepatic lipid metabolism, was the most significantly downregulated kinase in ALD-rat liver (p<0.05, FC>1.5). The upregulated kinases were associated with critical pathways, including PI3K-Akt, neurodegeneration, EGFR tyrosine kinase inhibitor resistance, cellular senescence, cAMP signaling, aldosterone secretion, mTOR, TNF-α, Th17 signaling, TGF-β, platelet activation, and others, with more than five kinases enriched in each pathway, detailed in Supplementary table-2.

Significant reductions were observed in kinases associated with phosphatidylinositol phosphate (PIP) synthesis at early and late endosome membranes, insulin secretion, circadian rhythm regulation, axon guidance, and others (Figure-3F, Supplementary table-2). Alterations in key signalling pathways, including FoxO, MAPK, and TLR4, were evident in ALD-rat liver tissues. Notably, 34 kinases and phosphoproteins associated with the MAPK pathway were significantly upregulated, while 8 kinases were downregulated (Figure-3G). These findings underscore the prominent role of MAPK signalling in driving inflammatory responses in ALD. KEGG pathway analysis identified TGFβR1, ROS production and LPS as potential upstream activators, linking fibrogenic (TGFβR1) and inflammatory (LPS) signalling in ALD pathogenesis.

Overall, temporal kinome analysis revealed that inflammatory kinases, particularly MAPK, drive ALD progression. Additionally, alcohol-induced liver damage was associated with neurodegeneration, senescence, and sodium imbalance, suggesting potential therapeutic targets for ALD management.

### Kinome profile of ALD-rat circulating monocytes

Systemic inflammation is a key feature of ALD. To investigate kinome alterations associated with this, circulating monocytes were analysed. Principal Component Analysis (PCA) revealed clear differences in the monocyte kinome profile of ALD-rats compared to controls (Figure-4A)

**Figure 4.**
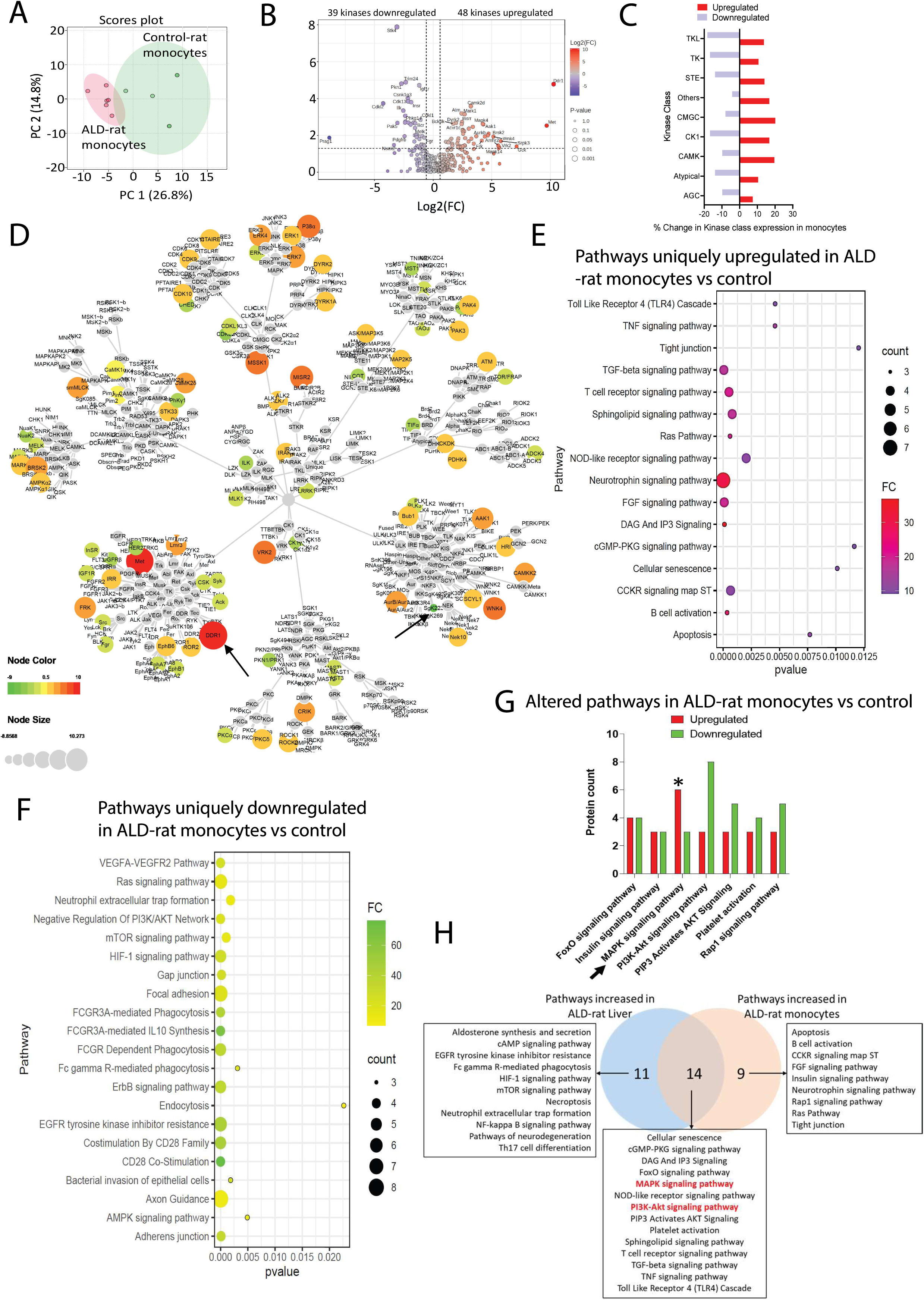
Kinome profile of ALD-rat circulating monocytes. A: Principle component analysis showing segregation in circulating monocytes kinome profile of ALD rats and control rats at 24 weeks. B: Volcano plot showing differential kinases in ALD monocytes compared to healthy rats. 48 kinases were significantly upregulated and 39 kinases were significantly downregulated in ALD rats. C: Bar plot showing percentage change in kinase class expression in ALD monocytes compared to healthy control monocytes. D: Kinome map representing differential kinases in ALD liver. Red= high expression, green: low expression. DDR1 had highest expression whereas PRAG1 (Skg223) had lowest expression in ALD monocytes compared to healthy monocytes. E: Dot plot showing pathways significantly upregulated in ALD monocytes compared to control. F: Dot plot showing pathways significantly downregulated in ALD monocytes compared to control. G: Bar plot showing altered pathways in ALD monocytes (involving both upregulated and downregulated kinases). 6 kinases associated with MAPK signaling were upregulated. H: Venn diagram showing pathways uniquely upregulated in ALD liver, aALD monocytes and commonly upregulated in ALD liver and monocytes.

A total of 345 kinases were identified of them 48 were significantly upregulated, and 39 were significantly downregulated in ALD-rat monocytes compared to healthy controls (p<0.05, FC> 1.5, Figure-4B). Class wise distinction of kinome profile (Figure-4C) along with a kinome tree map (Figure-4D), illustrated the differentially expressed kinases, with DDR1 showing the most pronounced upregulation, indicating a pro-inflammatory and migratory phenotype of ALD-rat monocytes (Figure-4D). Conversely, PRAG1 exhibited significant downregulation, suggesting insufficient regulation of pathways that counteract fibrosis and pro-inflammatory signaling, potentially leading to sustained inflammation [19] (Figure-4D). The most significantly upregulated kinases were associated with pathways such as TLR4 signaling, TNF signaling, TGF-β signaling, neurotrophin signaling, and sphingolipid signaling (p<0.05, FC>1.5, Figure-4E). In contrast, pathways linked to downregulated kinases included gap junctions, tight junctions, NET formation, phagocytosis, bacterial invasion, and IL-10 synthesis (Figure-4F). Notably, pathways such as FoxO signaling, insulin signaling, MAPK signaling (33 kinases up- and 8 kinases downregulated), and PI3K-Akt were dysregulated in ALD-rat monocytes (Figure-4G).

Venn analysis was performed to identify pathways commonly upregulated in both ALD-rat liver tissues and ALD-rat monocytes. The enriched pathways included cellular senescence, FoxO signaling, MAPK signaling, TLR4 signaling, TNF signaling, TGF-β signaling, and others (Figure-4H).

Together, our results highlight significant alterations in kinase expression in monocytes associated with ALD. The pronounced upregulation of pro-inflammatory kinases, such as DDR1, and MAPK14 alongside the downregulation of regulatory kinases like PRAG1, underscores critical pathways involved in inflammation and fibrosis. These findings suggest that targeting specific kinases may offer therapeutic potential for modulating inflammatory responses and reducing liver damage in the context of ALD.

### Temporal change in Kinome profile of ALD-rat liver and circulating monocytes

To investigate the progressive impact of alcohol on the pathophysiology and signalling pathways in ALD, liver tissues and circulating monocytes were collected at weeks 8, 12, 16, 20, and 24. Phosphoproteomic analysis was performed to characterize the temporal alterations in the kinome profile. Linear regression analysis identified 24 kinases that were upregulated and associated to cAMP signaling, endocrine resistance, IL-17 signaling, and TLR signaling whereas 31 kinases were downregulated over time in the liver (R²>0.85) (Figure-5A) and were linked to focal adhesion, tight junctions, and platelet activation, suggesting progressive disorganization of the extracellular matrix in liver due to decreased activity of ECM-maintenance kinases. Interestingly kinase analysis also identified some dysregulated pathways such as MAPK, insulin, PI3K-Akt, and Ras signaling pathways and others (Figure-5A).

**Figure 5.**
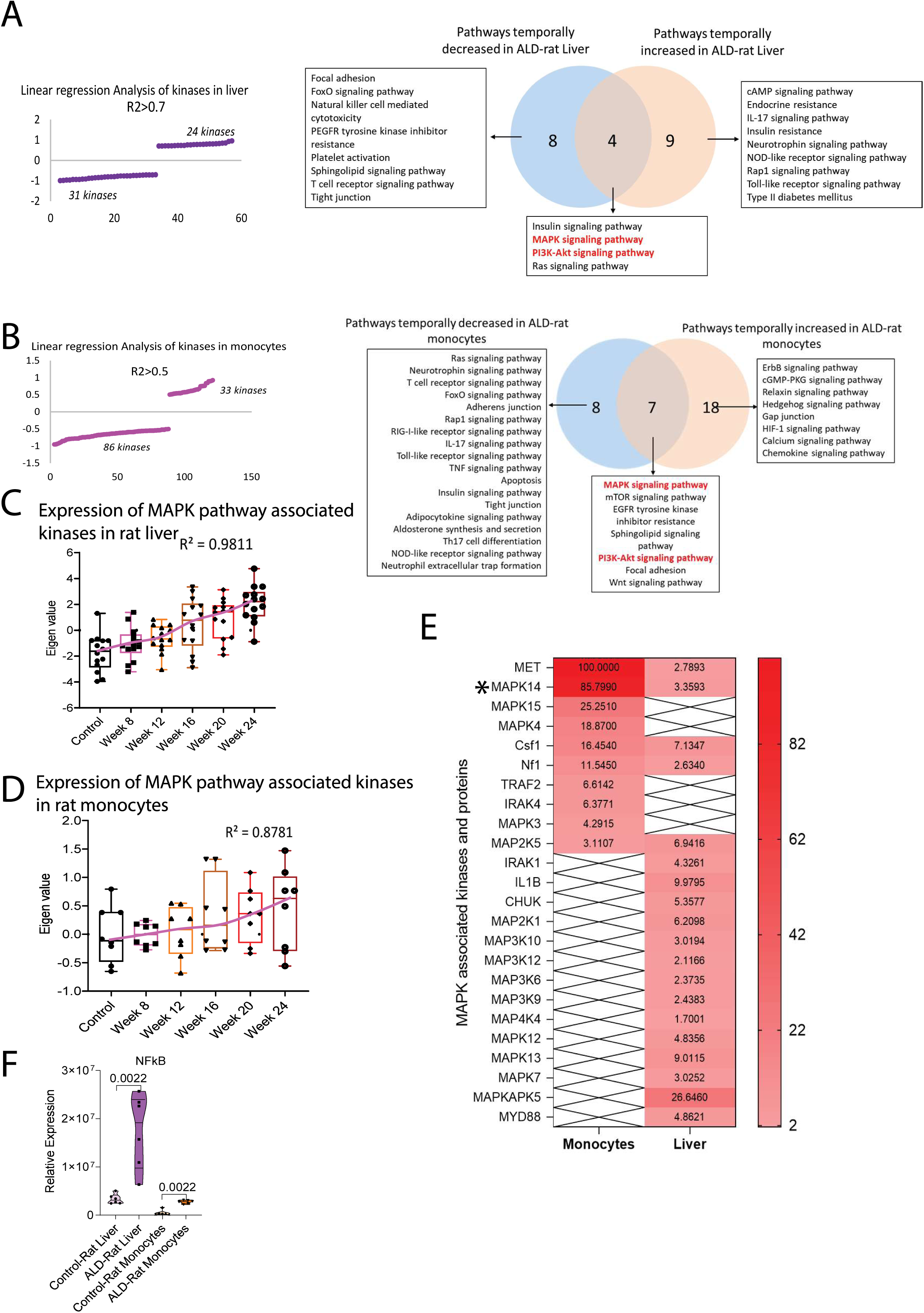
Temporal change in Kinome profile of ALD-rat liver and circulating monocytes. A: Linear regression analysis showing pathways associated with temporally increased 24 kinases and temporally decreased 31 kinases in ALD liver. B: Linear regression analysis showing pathways associated with temporally increased 24 kinases and temporally decreased 31 kinases in ALD monocytes. C: Whisker plot showing temporal increase in MAPK associated kinases in ALD liver over the time (R^2 0.981). D: Whisker plot showing temporal increase in MAPK associated kinases in ALD monocytes over the time (R^2 0.878). E: Expression of MAPK pathway associated kinases in ALD liver and monocytes. F: Expression of NFkB in ALD liver and monocytes compared to their respective controls.

Similarly in circulating monocytes, 33 kinases were temporally up-regulated, while 86 kinases were temporally downregulated (R²>0.5, Figure-5B). Upregulated kinases were enriched in pathways such as chemokine signaling and HIF-1 signaling, whereas downregulated kinases were associated with TCR signaling, neurotrophin signaling, IL-17 signaling, TNF signaling, and apoptosis. Gradual dysregulation was also noted in pathways such as MAPK, mTOR, and PI3K-Akt signaling (Figure-5B).

### Pinpointing Targetable Kinases in ALD rats

To identify potential therapeutic targets for controlling inflammation and its associated effects in ALD, pathways that were upregulated both in ALD-rat liver tissue and circulating monocytes, were examined. The MAPK pathway was consistently active in both ALD-rat liver and monocytes, showing a gradual increase over time (Figure-5C and Figure 5D). The expression patterns of kinases involved in this pathway are visualized in a heatmap (Figure-5E).

Pathway analysis of significantly upregulated phosphoproteins and kinases in ALD-rats, utilizing KEGG mapper, revealed a prominent activation of the MAPK pathway. Further investigation into this pathway identified potential upstream activators, namely TGFβR1, LPS and ROS production. These findings suggest a mechanistic link between fibrogenic signalling (TGFβR1) and inflammatory stimuli (LPS) in the pathogenesis of ALD (Supplementary Figure-3).

The MAPK pathway’s activation subsequently leads to the phosphorylation and activation of downstream kinases, most notably IKK (IκB Kinase) and p38 MAPK [12]. IKK activation is a critical step in the canonical NF-κB signalling pathway, leading to the translocation of NF-κB into the nucleus and the transcription of pro-inflammatory genes. Similarly, p38 MAPK activation also converges on NF-κB, either directly or through intermediate kinases, further amplifying the inflammatory response. On analysing the expression of NF-κB, we found significant increase in NFKB in both ALD-rat liver tissues and circulating monocytes (Figure-5F), confirming the sustained activation of this pathway and its central role in driving inflammation.

#### Validation: Modulation of NFkB activation using 70nM PS1145, 30nM PH797804 and 25uM Resveratrol

To mitigate inflammation in ALD, we targeted NF-κB, a key regulator of pro-inflammatory cytokines such as IL-6 and TNF-α, which contribute to liver damage [12]. Three inhibitors were selected for their ability to modulate NF-κB and MAPK signalling (Figure 6A): PS1145 (an IKKβ inhibitor that prevents IκB degradation, retaining NF-κB in the cytoplasm), PH797804 (a selective p38-MAPK inhibitor that disrupts TLR4-driven inflammatory cascades), and resveratrol (a polyphenol known to inhibit TLR4-mediated MAPK and NF-κB activation, serving as a positive control) and prednisolone. Their efficacy was assessed in THP-1 monocytes, HepG2 hepatocytes, PBMCs from healthy donors and SAH patients and in NIAAA mouse binge model (Figure-6A).

**Figure 6.**
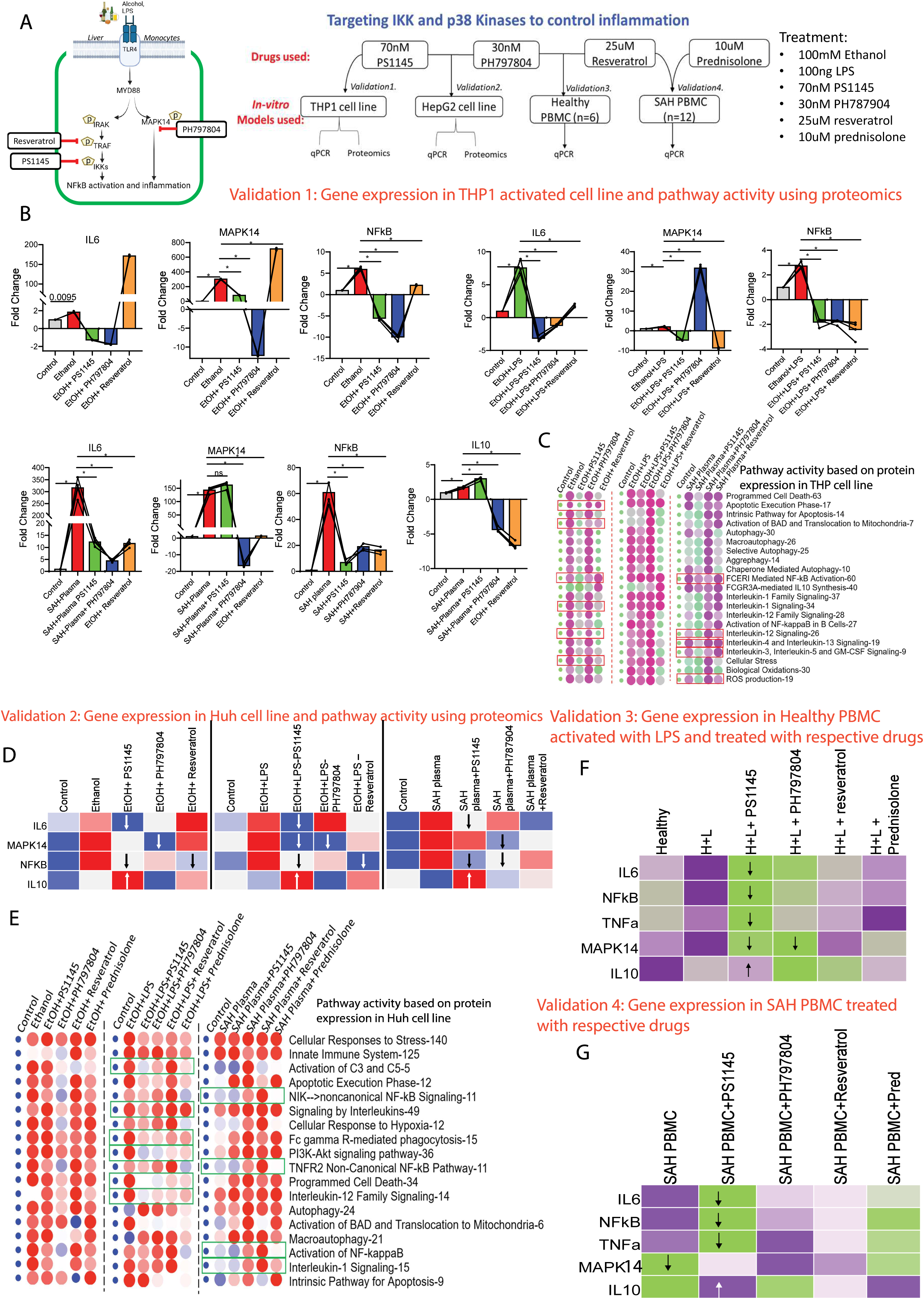
Pinpointing Targetable Kinases in ALD rats: A: Validation experiments. In alcoholic liver disease (ALD), activation of the MYD88 pathway leads to phosphorylation of IRAK, TRAF, and IKKs, resulting in IKK degradation and the release of NF-κB, which drives inflammation. The efficacy of PS1145, PH797804, and resveratrol was evaluated in controlling MYD88 pathway-induced inflammation using in vitro models, including THP-1 and HepG2 cell lines, as well as PBMCs from healthy individuals and patients with severe alcohol-associated hepatitis (SAH). B: Gene expression analysis of IL6, MAPK, NF-κB and IL10 in THP-1 cells treated with ethanol, ethanol + LPS, and patient plasma, followed by treatment with 70 nM PS1145, 30 nM PH797804, and 25 µM resveratrol. C: Heat map showing pathway activity of THP1 cell line treated with ethanol, ethanol + LPS, and patient plasma, followed by treatment with 70 nM PS1145, 30 nM PH797804, and 25 µM resveratrol. Red box highlights the pathway specifically downregulated by PS1145. D: Gene expression analysis of IL6, MAPK, NF-κB and IL10 in HepG2 cells treated with ethanol, ethanol + LPS, and patient plasma, followed by treatment with 70 nM PS1145, 30 nM PH797804, and 25 µM resveratrol. PS1145 stimulated IL10 production more efficiently compared to other drugs used. E: Heat map showing pathway activity of HepG2 cell line treated with ethanol, ethanol + LPS, and patient plasma, followed by treatment with 70 nM PS1145, 30 nM PH797804, 25 µM resveratrol and 10uM prednisolone. Green box highlights the pathway specifically downregulated by PS1145. F: Gene expression of IL6, NFkB, TNFa, MAPK14, and IL10 in healthy PBMC treated with LPS further treated with 70 nM PS1145, 30 nM PH797804, 25 µM resveratrol and 10uM prednisolone. G: Gene expression analysis of IL6, NF-κB, TNF-α, MAPK14, and IL10 in PBMCs from severe alcohol-associated hepatitis (SAH) patients treated with 70 nM PS1145, 30 nM PH797804, 25 µM resveratrol, and 10 µM prednisolone. PS1145 significantly reduced IL6, NF-κB, and TNF-α expression while enhancing IL10 production.

#### Validation Model 1: Evaluation of the efficacy of PS1145, PH797804, and resveratrol for amelioration of inflammation in the THP-1 cell line

Activated THP-1 monocytes were stimulated with 150 mM ethanol, 150 mM ethanol + 100 ng/mL LPS, and 10% SAH plasma respectively. These cells were subsequently treated with 70 nM PS1145, 30 nM PH797804, or 25 µM resveratrol for 24 hours and the expression of key inflammatory and anti-inflammatory markers were analysed as detailed in Supplementary Figure-4.

PS1145 demonstrated the most potent anti-inflammatory effects by significantly down-regulating IL-6 and NF-κB while up-regulating IL-10 (p<0.05). In contrast, PH797804 and resveratrol exhibited moderate anti-inflammatory effects but were less effective than PS1145 (p<0.05, Figure-6B). These findings highlight the therapeutic potential of PS1145 in suppressing inflammation in ALD-rats by directly targeting NF-κB activation and shifting the inflammatory balance towards an anti-inflammatory phenotype.

To further delineate the molecular mechanisms underlying PS1145-mediated NF-κB suppression, we performed untargeted proteomics. PS1145 effectively downregulated pathways associated with apoptosis, FCERI-mediated NF-κB activation, IL-12, IL-4, IL-13, oxidative stress, and reactive oxygen species (ROS) production (p<0.05, FC>1.5, Figure-6C).

#### Validation model 2: Evaluation of the efficacy of PS1145, PH797804, and resveratrol in the HepG2 cell line

Similarly, in HepG2 hepatocytes exposed to ethanol, ethanol + LPS, or 10% SAH plasma for 24 hours, treatment with PS1145, PH797804, resveratrol, and prednisolone revealed PS1145 as the most effective agent in suppressing IL-6 and NF-κB expression while stimulating IL-10 production (p<0.05, Figure-6D). Proteomic analysis of these samples demonstrated a significant down-regulation of complement activation (C3 and C5), non-canonical NF-κB signalling, TNFR2-mediated NF-κB activation, and IL-1 pathways, further confirming PS1145-mediated attenuation of inflammatory signalling (p<0.05, Figure-6E).

#### Validation model 3 and 4: Evaluation of the efficacy of PS1145, PH797804, and resveratrol in the primary PBMCs

To extend these findings to primary immune cells, we tested the efficacy of PS1145, PH797804, resveratrol, and prednisolone in PBMCs isolated from healthy individuals and SAH patients. Healthy PBMCs (n=6) were stimulated with 100 ng/mL LPS and subsequently treated with the respective inhibitors. Similar to previous observations, PS1145 significantly reduced the expression of IL-6, NF-κB, TNF-α, and MAPK14, while enhancing IL-10 production (Figure-6F). In SAH-derived PBMCs (n=12), PS1145 exhibited superior efficacy compared to PH797804, resveratrol, and prednisolone in suppressing IL-6, NF-κB, TNF-α, and MAPK14, while promoting IL-10 expression (Figure-6G). These results collectively indicate that inhibiting NF-κB through PS1145 effectively attenuates inflammatory cytokine production (better than PH797804 or resveratrol) while reinforcing anti-inflammatory pathways in ALD.

By suppressing NF-κB activation and promoting IL-10, PS1145 holds significant therapeutic promise in mitigating inflammation-driven liver pathology in ALD.

### PS1145 Ameliorates Alcohol-Induced Liver Injury via Dual Modulation of Inflammation and Metabolism in the NIAAA Model

To evaluate the therapeutic efficacy of PS1145 and PH797804 in alcoholic liver disease, the NIAAA chronic-binge ethanol mouse model was employed, with treatments administered as detailed in the methods (Figure 7A). Among the groups, prednisolone-treated mice exhibited the greatest change in body weight, followed by ALD controls, PS1145, and PH797804 (Figure 7B). The lowest liver-to-body weight ratio was observed in the PS1145 group, suggesting reduced hepatomegaly, while the most pronounced spleen atrophy occurred in ALD controls and prednisolone-treated mice (Figure 7B). Notably, prednisolone-treated mice also displayed physiological distress, including shivering, and showed the highest mortality rate, with one death before and one after the binge, whereas the ALD group had one death (Figure 7B). Histological analysis (H&E staining) demonstrated that PS1145 treatment significantly resolved hepatic steatosis and inflammation compared to prednisolone and PH797804, indicating that IKK inhibition not only suppresses NF-κB-driven inflammation but also improves hepatic lipid metabolism (Figure 7C). Proteomic analysis of liver tissues, followed by k-means clustering, identified several distinct clusters modulated by PS1145 (Figure 7D). The brown cluster, specifically downregulated by PS1145, was enriched for pathways such as tryptophan catabolism, neutrophil degranulation, MHC antigen presentation, mitochondrial fatty acid β-oxidation, and cytokine signalling, indicating effective suppression of inflammation and normalization of metabolic processes. Interestingly the expression of IL36R involved in dimerization of TLR receptor and functionality was specifically reduced by PS1145 treatment compared to other group (p<0.05, Figure 7F). This observation suggesting that PS1145 not only works through inhibition of phosphorylation of IKK but also compromise the TLR receptor assembly. In addition, the blue cluster, also downregulated by PS1145 compared to PH797804 and prednisolone, comprised proteins involved in cellular stress response, MAPK4/6 signalling, sulfide oxidation, and β-oxidation of very long-chain fatty acids, further supporting the reversal of alcohol-induced liver pathology (Figure 7E). In contrast, the light sky blue and blue-violet clusters, which were upregulated by PS1145, included proteins associated with GM-CSF signalling, methionine degradation, glutathione biosynthesis, regulation of pyruvate metabolism, carnitine synthesis, tryptophan degradation, carbohydrate metabolism, and biological oxidation (Figure 7E). These findings suggest that PS1145 not only attenuates hepatic inflammation (compromising TLR dimerization and also by inhibition of IKK phosphorylation) and steatosis but also restores metabolic and antioxidative balance. Collectively, these results highlight the superior therapeutic potential of PS1145 over PH797804 and prednisolone in the treatment of ALD.

**Figure 7:**
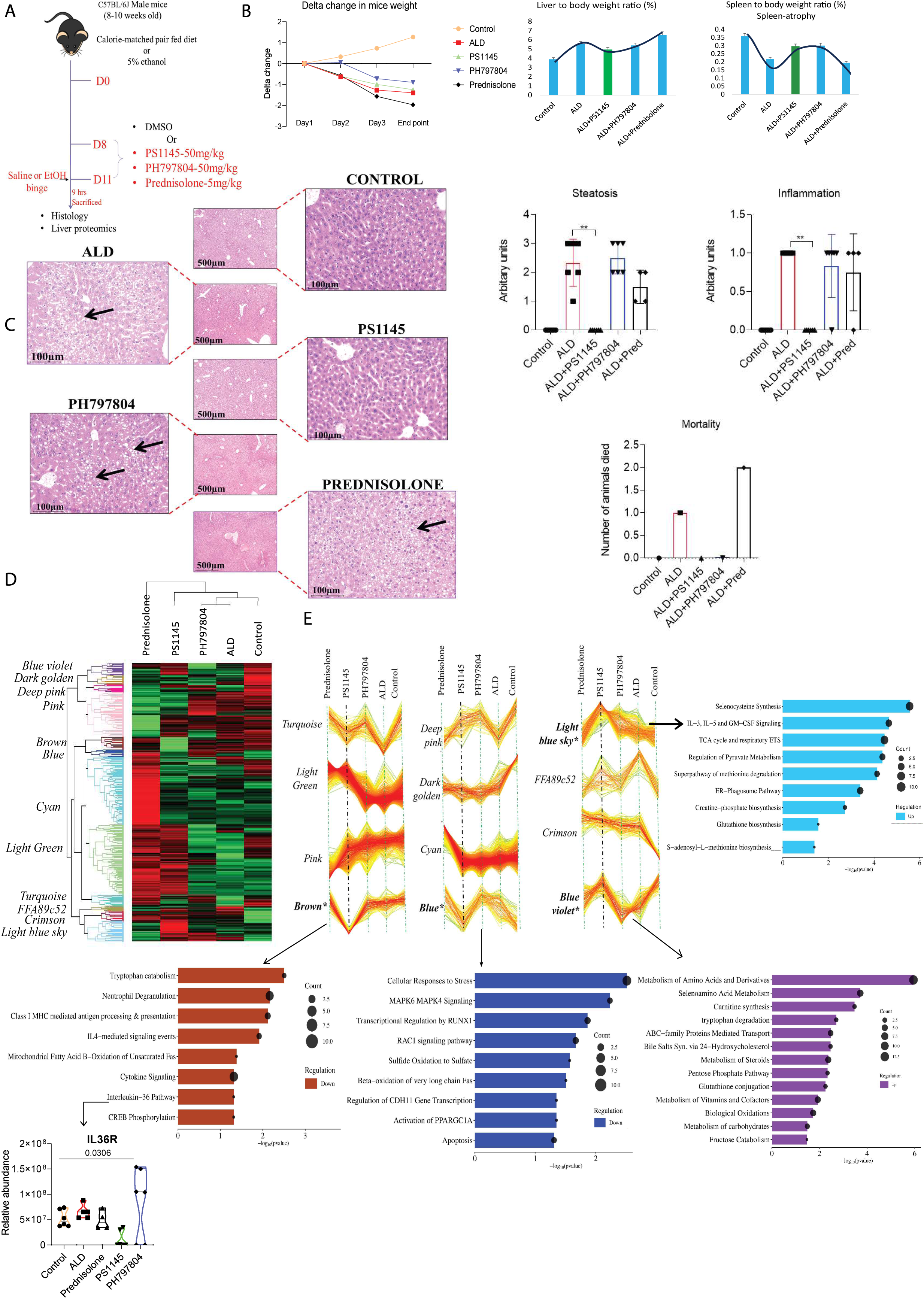
PS1145 Ameliorates Alcohol-Induced Liver Injury via Dual Modulation of Inflammation and Metabolism in the NIAAA Model: A: Schematic of NIAAA binge ethanol mouse model and treatment with respective therapeutic agents. B: Line graph depicting body weight changes over time; bar plots showing liver-to-body weight and spleen-to-body weight ratios at end time point. C: Representative liver histology sections showing steatosis and inflammation in ALD mice treated with respective drugs. Bar plots quantify steatosis score, inflammation score, and mortality rates across treatment groups. D: K-mean clustering heat map comparing NIAAA mice treated with respective drugs. E: 12 Clusters identified by K-mean clustering and pathways associated with brown, blue, light blue sky, and blue-violet cluster.

## DISCUSSION

Kinases are fundamental orchestrators of cellular signalling, regulating processes such as inflammation, apoptosis, and fibrosis, each of which is central to the pathogenesis of ALD [18] [3]. Inflammation plays a key role in disease progression, making its modulation a potential therapeutic target for mitigating ALD.

Through comprehensive kinome and phosphor-proteomic profiling, our study mapped the dynamic landscape of kinase activity in both liver tissue and circulating monocytes during ALD progression. This unbiased approach revealed profound dysregulation of key signalling networks, confirming the persistent upregulation of the MAPK and NF-κB pathways—particularly MAPK14 (p38) and IKK—as critical drivers of inflammation and hepatic injury in ALD.

Using the NIAAA chronic-binge ethanol mouse model and other *in-vitro* models, we evaluated the therapeutic potential of two targeted kinase inhibitors, PS1145 (IKK inhibitor) and PH797804 (p38 MAPK inhibitor), in direct comparison with the standard-of-care agent prednisolone. Our results demonstrated that PS1145 provided superior protection against ethanol-induced liver injury, as evidenced by reduced liver-to-body weight ratios, improved histological features (notably the resolution of steatosis and inflammation), and lower mortality rates. Proteomic clustering further revealed that PS1145 uniquely downregulated inflammatory and metabolic stress pathways, while upregulating cytoprotective and metabolic restoration pathways. This dual modulation suggests that PS1145 not only suppresses pro-inflammatory signaling via IKK/NF-κB inhibition and via inhibiting the dimerization of TLR receptor (decrease expression of IL-36R) but also promotes metabolic and antioxidative homeostasis, features not observed with PH797804 or prednisolone.

Recent studies have established that TLR4-driven inflammation dominates both liver and monocytes in ALD, with key kinases: IKK, MAPK14, TRAF, and IRAK highly active and implicated in systemic inflammation [20]. To target this pathway, we tested PS1145 (IKK inhibitor), PH797804 (MAPK14 inhibitor), and resveratrol (TRAF inhibitor), with prednisolone as a standard-of-care comparator.

To our knowledge, this is the first study directly comparing PS1145 and PH797804 in ALD. We demonstrate PS1145’s novel anti-inflammatory effect via IKK phosphorylation inhibition and reduced IL-36R expression (TLR dimerization), highlighting its unique therapeutic potential in alcoholic liver disease.

Kinome analysis led to the identification of 497 and 345-kinases in liver tissue and monocytes respectively which provides a foundational resource for understanding kinase-driven regulatory networks in ALD. The observed subcellular localization of kinases across different compartments, including cytoplasm, nucleus, plasma membrane, Golgi apparatus, endoplasmic reticulum, and mitochondria, aligns with existing literature on compartment-specific signalling dynamics in ALD patients [21].

ALD progression is marked by profound alterations in kinase expression, as evidenced by PCA showing distinct clustering of ALD and control samples. The widespread dysregulation of kinases in ALD suggests significant perturbations in hepatic signalling pathways, contributing to disease progression. The increased expression of TGFBR1, a key mediator of fibrosis, underscores the activation of TGF-β signalling, which drives extracellular matrix remodelling and fibrogenesis [22]. Additionally, increase in kinases such as GUCY2D, PHKG2, and CAMK1D, known to regulate hepatocellular stress responses, metabolic homeostasis, and inflammatory pathways, highlight the multifaceted impact of alcohol-induced stress on liver. Concordant activation of these kinases suggests a convergence of fibrotic, metabolic, and inflammatory mechanisms that exacerbate ALD pathophysiology.

Conversely, the downregulation of kinases such as SBK2 indicates disrupted metabolic control that may contribute to hepatic steatosis and dysfunction [23]. The enrichment of dysregulated kinases in pathways such as PI3K-Akt, mTOR, TNF-α, and TGF-β signalling reinforces their involvement in hepatocyte survival, inflammation, and fibrotic remodelling [24]. The suppression of kinases involved in insulin signalling suggests potential impairments in glucose metabolism and hepatic insulin resistance ALD [25]. These findings highlight the intricate interplay between kinase dysregulation, metabolic dysfunction, and inflammatory signalling in ALD progression.

Circulating monocytes had specific increase of pro-inflammatory kinases such as DDR1 and MAPK14, while PRAG1 was notably downregulated. The role of DDR1 in promoting monocyte migration and tissue infiltration has been previously reported in inflammatory liver disease [26]. The activation of MAPK14 is particularly relevant, given its well-established role in TNF-α and IL-6 production (key cytokines driving ALD pathology) [27].

Longitudinal kinome analysis in ALD liver and monocyte highlights the progressive dysregulation of kinases over the course of ALD development. The sustained activation of kinases linked to cAMP, IL-17, and TLR signalling suggests a reinforcing cycle of inflammatory and fibrotic responses. Meanwhile, the gradual suppression of kinases involved in focal adhesion and tight junction maintenance suggests a progressive loss of hepatocyte integrity, potentially leading to increased hepatocyte detachment and extracellular matrix destabilization [28].

Our study confirmed MAPK pathway as a consistently upregulated signalling cascade in both liver tissue and circulating monocytes of rodents with experimental ALD, suggesting its crucial role in systemic inflammation and disease progression. We observed a temporal upregulation of this pathway, with the most prominent kinases being TGFBR1 along with significant increase in the LPS and ROS, indicating that ROS induced inflammation drives the fibrogenesis in ALD (Figure-5C-E, and Supplementary Figure-4).

Dysregulation of the MAPK pathway has been implicated in various liver diseases, including ALD [29]. Elevated NF-κB expression in both liver tissues and circulating monocytes (Figure-5F) further underscores the sustained activation of inflammatory pathways in ALD as it is central regulator of pro-inflammatory cytokine production, including IL-6 and TNF-α [9, 12]. Therefore, modulating this pathway in ALD could be beneficial.

NFkB was targeted using three pharmacological inhibitors PS1145, PH797804, and resveratrol, and prednisolone as comparator each acting on distinct components of the NF-κB and MAPK signalling. PS1145, an IKKβ inhibitor, is a well-characterized compound known to block the phosphorylation and subsequent degradation of IκB, effectively sequestering NF-κB in the cytoplasm [30]. This mechanism not only suppresses pro-inflammatory transcriptional activity but tunes the cascade in a manner that it promotes the expression of anti-inflammatory genes, such as IL-10, contributing to a shift in the inflammatory balance [31].

While PH797804, a selective p38 MAPK inhibitor, targets a different point in the inflammatory cascade. The rationale for including resveratrol, a polyphenolic compound, stems from its well-established anti-inflammatory properties, specifically its ability to inhibit TLR4-mediated MAPK signaling and has also been used in ALD treatment. The comparison of the pharmacological inhibitors was carried out across distinct cellular models relevant to ALD depicted in Figure-6A.

PS1145 demonstrated superior efficacy in suppressing IL-6 and NF-κB while upregulating IL-10 in THP-1 monocytes (Figure-6B). It’s worth noting that studies have also found IKK inhibition to be effective in animal models of inflammatory bowel disease (IBD), thereby suggesting the systemic anti-inflammatory properties of IKK inhibition (M Pasparakis et. al 2008). The less pronounced effects of PH797804 and resveratrol are consistent with other studies demonstrating the importance of IKKβ as a central node in NF-κB activation, though their effects still point to the impact of MAPK and TLR4 on inflammatory processes.

Proteomic analysis revealed, PS1145 effectively downregulated apoptosis, FCERI-mediated NF-κB activation, IL-12, IL-4, IL-13, oxidative stress, and reactive oxygen species (ROS) production (Figure-6C). These findings suggest a broad mechanism of action for PS1145, extending beyond direct NF-κB inhibition. FCERI activation is known to trigger mast cell degranulation and the release of inflammatory mediators, and the suppression of this pathway further underscores the anti-inflammatory effects of PS1145. The downregulation of ROS production is particularly relevant, given the role of oxidative stress in ALD pathogenesis. PS1145 demonstrated similar efficacy in suppressing IL-6 and NF-κB expression while stimulating IL-10 production in ethanol, ethanol + LPS, or SAH plasma-exposed HepG2 hepatocytes (Figure-6D). Proteomic analysis also confirmed the attenuation of inflammatory signalling mediated by complement activation (C3 and C5), non-canonical NF-κB signalling, TNFR2-mediated NF-κB activation, and IL-1 pathways (Figure-6E). TNFR2-mediated NF-κB activation has been implicated in chronic inflammatory diseases, and its suppression by PS1145 further highlights the therapeutic potential of this compound. The C3 and C5 activation are usually associated with inflammation mediated cell death [32].

To validate the clinical relevance of these findings, we tested the efficacy of the pharmacological agents in healthy and SAH PBMCs as well. Similar to THP-1 monocytes and HepG2 hepatocytes, PS1145 exhibited superior efficacy compared to PH797804, resveratrol, and prednisolone in suppressing IL-6, NF-κB, TNF-α, and MAPK14, while promoting IL-10 expression. These results collectively indicate that inhibiting NF-κB through PS1145 effectively attenuates inflammatory cytokine production while reinforcing anti-inflammatory pathways in ALD.

By suppressing NF-κB activation and promoting IL-10, PS1145 holds significant therapeutic promise in mitigating inflammation-driven liver pathology in ALD. Future studies should explore the clinical translatability of these findings. This will not only help to develop targeted therapeutic interventions but also pave the way to identify novel therapeutic targets.

Finally, using the NIAAA ethanol mouse model and in vitro systems, we compared PS1145 (IKK inhibitor), PH797804 (p38 MAPK inhibitor), and prednisolone. PS1145 showed superior hepatoprotection, reducing steatosis, inflammation, and mortality. Proteomic profiling revealed PS1145 uniquely downregulated inflammatory and metabolic stress pathways while enhancing cytoprotective responses, suggesting dual action IKK/NF-κB and TLR/IL-36R inhibition alongside restoration of metabolic and antioxidant balance, not seen with PH797804 or prednisolone.

There are certain limitations of this study. The use of a rodent model may not fully replicate human disease. Kinase activity was inferred from phosphoproteomics without direct functional validation. The therapeutic potential of kinase inhibitors requires further in vivo and clinical validation. Additionally, the study focuses on specific pathways, and broader systemic interactions in ALD remain unexplored.

## CONCLUSION

PS1145 holds significant therapeutic promise for mitigating inflammation-driven liver pathology in ALD by effectively attenuating inflammatory cytokine production (inhibition of phosporylation of IKK and TLR/IL-36R inhibition, Graphical abstract) and reinforcing anti-inflammatory pathways. Future studies should prioritize exploring the clinical translatability of these findings through in vivo studies and clinical trials, with the goal of developing targeted therapeutic interventions for ALD and identifying novel kinase-based therapeutic targets.

## Disclosure

All authors have declared no conflict of interest.

## Financial Support

The man power was supported by CSIR. The work was supported by ICMR (IIRP-2023-1575).

## Authors’ contribution

SKS, JSM, AG and MY conceptualized the work. YM, SY, VS, YM and NS, KY were responsible for sample processing and experimental work and were helped by NS, VB, SP, GT, RF, and BM. Veterinary help was provided by AK. Data analysis was performed by MY under the guidance of and JSM. The manuscript was drafted by MY, SKS, and JSM. All authors have approved this manuscript.

## Abbreviations

ALD: Alcohol-related Liver Disease
SAH: Severe alcohol-related hepatitis
AST: Aspartate aminotransferase
ALT: Alanine aminotransferase
ROS: Reactive Oxygen Species
LDC: Lieber–De Carli
LE: Long Evan
mTOR: Mammalian Target of Rapamycin
MAPK14: Mitogen-Activated Protein Kinase 14 or p38
NFkB: Nuclear Factor-kappa B
IL6: Interleukin-6
TNFa: Tumor Necrosis Factor alpha
IL10: Interleukin-10
FC: Fold Change

---

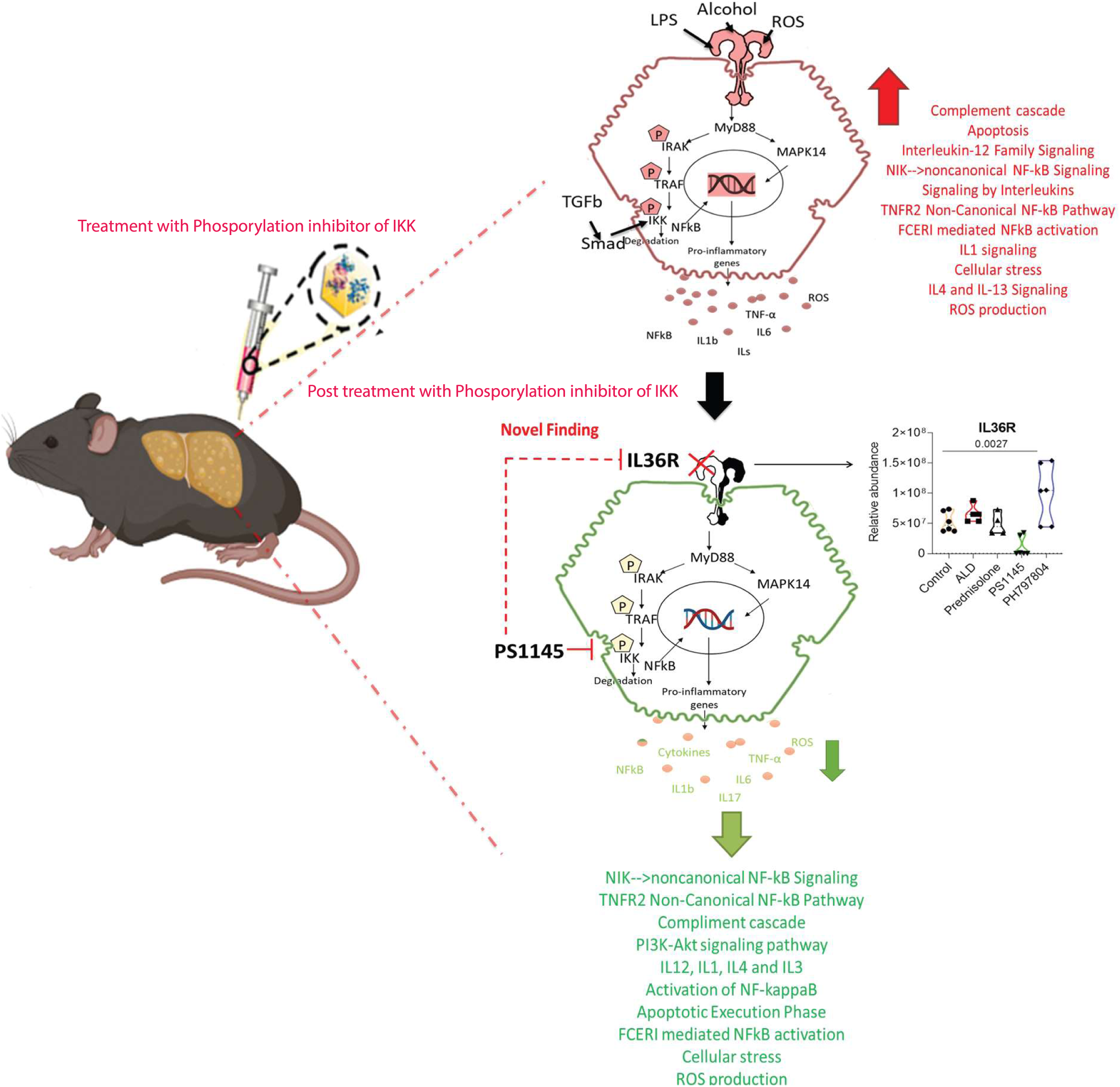

## Notes

### Competing Interest Statement

The authors have declared no competing interest.

## Reference

[1] Mackowiak B, Fu Y, Maccioni L, et al. Alcohol-associated liver disease. J Clin Invest 2024;134.

[2] Nagy LE. The Role of Innate Immunity in Alcoholic Liver Disease. Alcohol Res 2015;37:237–250.

[3] Koyama Y, Brenner DA. Liver inflammation and fibrosis. J Clin Invest 2017;127:55–64.

[4] Gao B, Bataller R. Alcoholic liver disease: pathogenesis and new therapeutic targets. Gastroenterology 2011;141:1572–1585.

[5] Arab JP, Diaz LA, Baeza N, et al. Identification of optimal therapeutic window for steroid use in severe alcohol-associated hepatitis: A worldwide study. J Hepatol 2021;75:1026–1033.

[6] Liu SY, Tsai IT, Hsu YC. Alcohol-Related Liver Disease: Basic Mechanisms and Clinical Perspectives. Int J Mol Sci 2021;22.

[7] Morales-Ibanez O, Affo S, Rodrigo-Torres D, et al. Kinase analysis in alcoholic hepatitis identifies p90RSK as a potential mediator of liver fibrogenesis. Gut 2016;65:840–851.

[8] Mandrekar P, Szabo G. Signalling pathways in alcohol-induced liver inflammation. J Hepatol 2009;50:1258–1266.

[9] Nowak AJ, Relja B. The Impact of Acute or Chronic Alcohol Intake on the NF-kappaB Signaling Pathway in Alcohol-Related Liver Disease. Int J Mol Sci 2020;21.

[10] Wu X, Liu YK, Iliuk AB, et al. Mass spectrometry-based phosphoproteomics in clinical applications. Trends Analyt Chem 2023;163.

[11] Savage SR, Zhang B. Using phosphoproteomics data to understand cellular signaling: a comprehensive guide to bioinformatics resources. Clin Proteomics 2020;17:27.

[12] Mulero MC, Huxford T, Ghosh G. NF-kappaB, IkappaB, and IKK: Integral Components of Immune System Signaling. Adv Exp Med Biol 2019;1172:207–226.

[13] Canovas B, Nebreda AR. Diversity and versatility of p38 kinase signalling in health and disease. Nat Rev Mol Cell Biol 2021;22:346–366.

[14] Tripathi G, Sharma N, Bindal V, et al. Protocol for global proteome, virome, and metaproteome profiling of respiratory specimen (VTM) in COVID-19 patient by LC-MS/MS-based analysis. STAR protocols 2022;3:101045.

[15] Yu LR, Veenstra T. Phosphopeptide enrichment using offline titanium dioxide columns for phosphoproteomics. Methods in molecular biology 2013;1002:93–103.

[16] Bujanda L, Garcia-Barcina M, Gutierrez-de Juan V, et al. Effect of resveratrol on alcohol-induced mortality and liver lesions in mice. BMC Gastroenterol 2006;6:35.

[17] Bertola A, Mathews S, Ki SH, et al. Mouse model of chronic and binge ethanol feeding (the NIAAA model). Nature protocols 2013;8:627–637.

[18] Bononi A, Agnoletto C, De Marchi E, et al. Protein kinases and phosphatases in the control of cell fate. Enzyme Res 2011;2011:329098.

[19] Wynn TA, Ramalingam TR. Mechanisms of fibrosis: therapeutic translation for fibrotic disease. Nat Med 2012;18:1028–1040.

[20] Mandrekar P. Signaling mechanisms in alcoholic liver injury: role of transcription factors, kinases and heat shock proteins. World journal of gastroenterology 2007;13:4979–4985.

[21] Aebersold R, Mann M. Mass-spectrometric exploration of proteome structure and function. Nature 2016;537:347–355.

[22] Dooley S, ten Dijke P. TGF-beta in progression of liver disease. Cell and tissue research 2012;347:245–256.

[23] Schroeder F, Atshaves BP, McIntosh AL, et al. Sterol carrier protein-2: new roles in regulating lipid rafts and signaling. Biochim Biophys Acta 2007;1771:700–718.

[24] Roy T, Boateng ST, Uddin MB, et al. The PI3K-Akt-mTOR and Associated Signaling Pathways as Molecular Drivers of Immune-Mediated Inflammatory Skin Diseases: Update on Therapeutic Strategy Using Natural and Synthetic Compounds. Cells 2023;12.

[25] Cheng Q, Li YW, Yang CF, et al. Ethanol-Induced Hepatic Insulin Resistance is Ameliorated by Methyl Ferulic Acid Through the PI3K/AKT Signaling Pathway. Front Pharmacol 2019;10:949.

[26] Bansod S, Saifi MA, Godugu C. Inhibition of discoidin domain receptors by imatinib prevented pancreatic fibrosis demonstrated in experimental chronic pancreatitis model. Sci Rep 2021;11:12894.

[27] Lo U, Selvaraj V, Plane JM, et al. p38alpha (MAPK14) critically regulates the immunological response and the production of specific cytokines and chemokines in astrocytes. Sci Rep 2014;4:7405.

[28] Rao R. Endotoxemia and gut barrier dysfunction in alcoholic liver disease. Hepatology 2009;50:638–644.

[29] Aroor AR, Shukla SD. MAP kinase signaling in diverse effects of ethanol. Life Sci 2004;74:2339–2364.

[30] Lung HL, Kan R, Chau WY, et al. The anti-tumor function of the IKK inhibitor PS1145 and high levels of p65 and KLF4 are associated with the drug resistance in nasopharyngeal carcinoma cells. Sci Rep 2019;9:12064.

[31] Schottelius AJ, Mayo MW, Sartor RB, et al. Interleukin-10 signaling blocks inhibitor of kappaB kinase activity and nuclear factor kappaB DNA binding. J Biol Chem 1999;274:31868–31874.

[32] Santiesteban-Lores LE, Carneiro MC, Isaac L, et al. Complement System in Alcohol-Associated Liver Disease. Immunol Lett 2021;236:37–50.

