## Supplementary methods for "Liver Kinome Profiling Identifies PS1145 as a Potential Therapeutic molecule for amelioration of Systemic Inflammation in Alcohol-related Liver Disease"

***ALD rat model development:***

The study involved the administration of ethanol to 6- to 8-week-old male and female Long Evans rats sourced from the National Brain Research Centre, India. Upon arrival, the rats were acclimatized to individual housing conditions maintained at 20 °C with 45% relative humidity and a 12-hour light/dark cycle, with unrestricted access to water and food for one week. Following this adaptation period, the rats were shifted from their standard chow diet to a Lieber-De Carli (LDC) control liquid diet (Catalog# F1259 BioServ, Frenchtown, NJ), in accordance with (14). The rats were then randomly divided into two groups: a control group (n=6) and an Alcohol-related liver Disease (ALD) group (n=6). The ALD group underwent a gradual introduction to ethanol over seven days, starting at 5% and increasing to 40%, continuing for a total of 24 weeks. The control group was pair-fed with the LDC diet to match the caloric intake from ethanol.

***Sample Collection:*** The rats were euthanized at various intervals: 8 weeks (n=5), 12 weeks (n=5), 16 weeks (n=3), 20 weeks (n=5), and 24 weeks (n=6). They were assessed for histological changes indicative of steatosis, inflammation, and fibrosis, alongside biochemical evaluations including liver function tests and lipid profiles.

Liver tissue processing followed established protocols as described by Sadeghipour and Babaheidarian (2019). The tissues were initially fixed in 10% formalin and subsequently immersed in isopropanol. To eliminate residual formalin, the tissues underwent a series of increasing alcohol concentrations before being cleaned with xylene. Finally, the tissues were embedded in wax to enhance structural integrity.

***Haematoxylin and Eosin Staining:***

Histological examination was conducted using Fischer's method. Tissue sections were rehydrated, stained with Mayer's hematoxylin for 30 seconds, rinsed in water for one minute, then stained with eosin for 10 to 30 seconds. After dehydration through graded alcohols, the slides were washed in xylene and prepared for microscopic examination using a Zeiss Axiophot microscope with varying objectives.

***Picro Sirius Red Staining:***

The Picro Sirius Red staining followed López's method. Paraffin-embedded sections underwent deparaffinization and rehydration through xylene and alcohol rinses. They were incubated in a solution of 0.1% Sirius Red mixed with saturated aqueous picric acid for one hour, then rinsed in acidified water containing hydrogen chloride before dehydration and mounting with DPX. This staining highlighted collagen in red while non-collagen components appeared orange. The stained sections were also examined under a Zeiss Axiophot microscope equipped with multiple objectives.

***Oil Red Staining:***

Frozen tissue sections were prepared and incubated with a freshly prepared Oil Red O solution to visualize neutral lipids and triglycerides. Following incubation, sections were rinsed and counterstained with hematoxylin to enhance contrast. The stained sections were examined under a microscope to assess lipid accumulation and morphology, highlighting lipid deposits within the tissues.

***Quantitative Real­Time PCR Analysis***

The measurement of gene expression via Real­Time PCR was performed as previously described. In brief, total RNA was extracted from liver tissues and cell lysate using the manual trizol method. Total RNA was read on a NanoDrop 2000 spectrophotometer (Thermo Fisher Scientific, Wilmington, DE), and cDNA was synthesized using RevertAid First Strand cDNA Synthesis Kit (#K1621). PCR amplification of the cDNA was performed by quantitative real­time PCR (qt-RT-PCR) using Powerup SYBR Green qPCR masterMix (Catalog number: A25742, Thermo Scientific™, USA)). The thermocycling protocol consisted of 2 minutes at 50°C, 10’ at 95°C, 40 cycles of 15 seconds at 95°C, 60 seconds at 60°C, and based on primer size 0 to 30 seconds at 72°C and finished with a melting curve ranging from 60–95°C to allow distinction of specific products.

**Proteomics sample preparation and run**:

50 µg equivalent protein was isolated from liver tissue lysate samples and were subjected to reduction with 10mM DTT at 60°C for 1Hr, alkylation with 10mM Iodoacetamide (IAA) in the dark at room temperature, and digestion for 24 hrs at 37°C using trypsin (Promega: V5280). Samples were desalted with C18 spin columns (PierceTM: 89870) and lyophilized. Finally, the samples were reconstituted in 0.1% formic acid. The peptides underwent nano-electrospray ionization followed by tandem mass spectrometry (MS/MS) using a Q-ExactiveTM Plus instrument (Thermo Fisher Scientific, San Jose, CA, United States). After enrichment on a trap column (75 μm x 2 cm, 3 μm, 100Å, nano Viper 2Pk C18 Acclaim PepMapTM 100) at a flow rate of 8 μl/min, they were separated on an analytical column (75 μm × 25 cm, 2 μm, 100Å, nano Viper C18, Acclaim PepMapTM RSLC). Peptides were eluted over a 120-minute gradient (3–95% buffer B: 80% acetonitrile in 0.1% formic acid) at a continuous flow rate of 800 nL/min. Mass spectrometry was performed using collision-induced dissociation and an electrospray voltage of 2.3 kV. Orbitrap analysis included full scan MS spectra at a resolution of 70,000 over the m/z range 350–1800. Protein identification employed the Mascot algorithm (Mascot 2.4, Matrix Science), with significance determined at p < 0.05 and q values (false discovery rate) also set at p < 0.05, maintaining a false discovery rate threshold of 0.01

***Activated monocytes isolation:***

Stored PBMCs were thawed, washed in PBS, and centrifuged at 400g for 10 minutes. The pellet was resuspended in RPMI-1640 with 10% FBS and 1% penicillin-streptomycin, then incubated at 37°C with 5% CO₂ for 4 hours. Non-adherent cells were removed, and adherent monocytes were lysed with RIPA buffer on ice for 1 hour. The lysate was collected, centrifuged at 10,000 rpm for 10 minutes, and the supernatant retained for phosphoproteomics analysis.

***THP1 and HepG2 cell line culture***

THP-1 cells, a human monocytic cell line, were differentiated into M0 macrophages using a standard PMA-induced activation protocol. The cells were initially cultured in RPMI-1640 medium supplemented with 10% fetal bovine serum (FBS) and 1% penicillin-streptomycin in a humidified incubator at 37°C with 5% CO₂. THP-1 cells in the exponential growth phase were seeded in tissue culture plates at a density of 1 × 10⁶ cells/mL. Phorbol 12-myristate 13-acetate (PMA) at a concentration 25ng/ml was added to the culture medium induce differentiation. The cells were incubated with PMA for 48 hours, during which they adhered to the culture plate surface, signifying their differentiation into macrophage-like cells. Following the incubation period, the PMA-containing medium was carefully removed, and the cells were washed with phosphate-buffered saline (PBS) to remove any residual PMA. The macrophages were then cultured in fresh RPMI-1640 medium without PMA for an additional 24 hours to allow them to stabilize as M0 macrophages.

HepG2 cell line was revived in DMEM media and was grown till 80% confluency and seeded in 6 well plate to conduct the experiment.

***Determining the Tolerated Dose of PS1145, PH787904, and Resveratrol in THP-1 Differentiated Cells Using MTT Assay***

The MTT assay was performed to determine the tolerated doses of PS1145, PH787904, and resveratrol in differentiated THP-1 macrophages. After the stabilization of the differentiated cells, PS1145, PH787904, and resveratrol were added at varying concentrations: PS1145 at 40 nM, 80 nM, 160 nM, and 320 nM; PH787904 at 15 nM, 30 nM, 60 nM, 120 nM, and 240 nM; and resveratrol at 25 µM, 50 µM, 100 µM, and 150 µM. The cells were treated with these concentrations for 24 hours.

Following the treatment period, 10 µL of MTT reagent (5 mg/mL in PBS) was added to each well and incubated for 4 hours at 37°C in a humidified incubator to allow the formation of formazan crystals. After incubation, the medium was carefully removed without disturbing the crystals, and 100 µL of dimethyl sulfoxide (DMSO) was added to each well to dissolve the formazan. The plate was gently agitated to ensure complete dissolution of the crystals.

The absorbance of each well was measured at 570 nm using a microplate reader, with a reference wavelength of 630 nm to account for background. The absorbance values were used to calculate cell viability as a percentage of untreated control cells. The results were analyzed to identify the maximum concentrations of PS1145, PH787904, and resveratrol that were tolerated by the cells without significant loss of viability.

***Evaluation of Anti-Inflammatory Effects of PS1145, PH787904, and Resveratrol Using THP-1 Macrophages:***

The experiment aimed to evaluate the efficacy of PS1145, PH787904, and resveratrol in controlling inflammation in THP-1 cells activated by different inflammatory stimuli. THP-1 cells, differentiated into macrophages, were seeded in appropriate culture plates and activated using 150 mM ethanol, 100 ng/mL lipopolysaccharide (LPS), a combination of 100 mM ethanol and 100 ng/mL LPS, or patient plasma for 24 hours. Untreated controls corresponding to each condition were also maintained. After the activation period, the cells were treated with 70 nM PS1145, 30 nM PH787904, or 25 µM resveratrol for an additional 24 hours.

Following the treatment, RNA was extracted for gene expression analysis using real-time PCR (RT-PCR) with primers for IL6, NFκB, MAPK, and IL10 in SAH plasma treated group, normalized to a housekeeping gene and analyzed using the ΔΔCt method. For proteomics, cell lysates were prepared, protein concentration quantified, and equal amounts digested for mass spectrometry analysis to identify differentially expressed inflammatory pathways.

The results from RT-PCR and proteomics were compared between treated and untreated cells under each activation condition to evaluate the efficacy of PS1145, PH787904, and resveratrol in modulating inflammatory responses. This approach provided insights into the compounds' ability to control inflammation at both transcriptional and protein levels.

***HepG2 cell line culture and treatment with PS1145, PH797804, and Resveratrol.***
