## Supplementary figures for "Liver Kinome Profiling Identifies PS1145 as a Potential Therapeutic molecule for amelioration of Systemic Inflammation in Alcohol-related Liver Disease"

**Supplementary figure 1:**

Table showing cellular component analysis with number of kinases enriched from specific cellular component in Liver

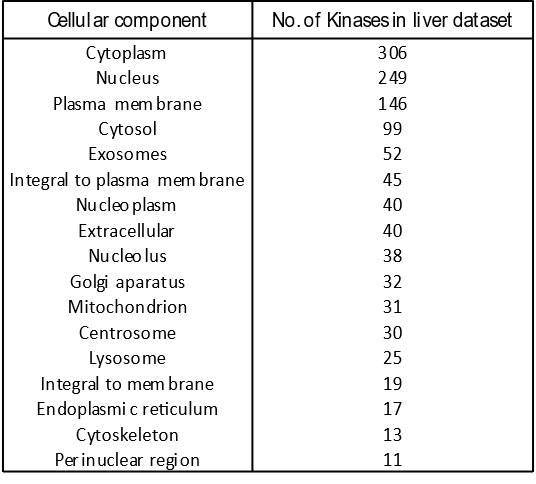

**Supplementary figure 2:**

Table showing cellular component analysis with number of kinases enriched from specific cellular component in monocytes

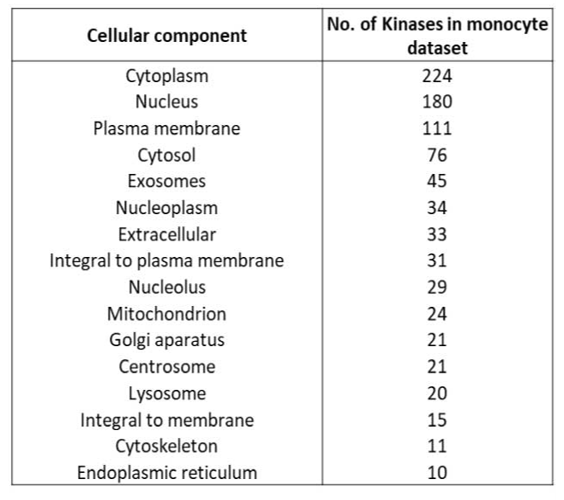

**Supplementary figure 3:**

In detail representation of MAPK Pathway in liver and monocytes. Highlighted kinases in orange yellow and blue colour shows the kinases found to be upregulated in ALD liver, monocytes and both in liver and monocytes respectively. A significant increase in classical pathways was seen in ALD. In addition, there was an increase in p38 kinase pathway by ROS production, IL1, TGFBR1, cytokines and others.

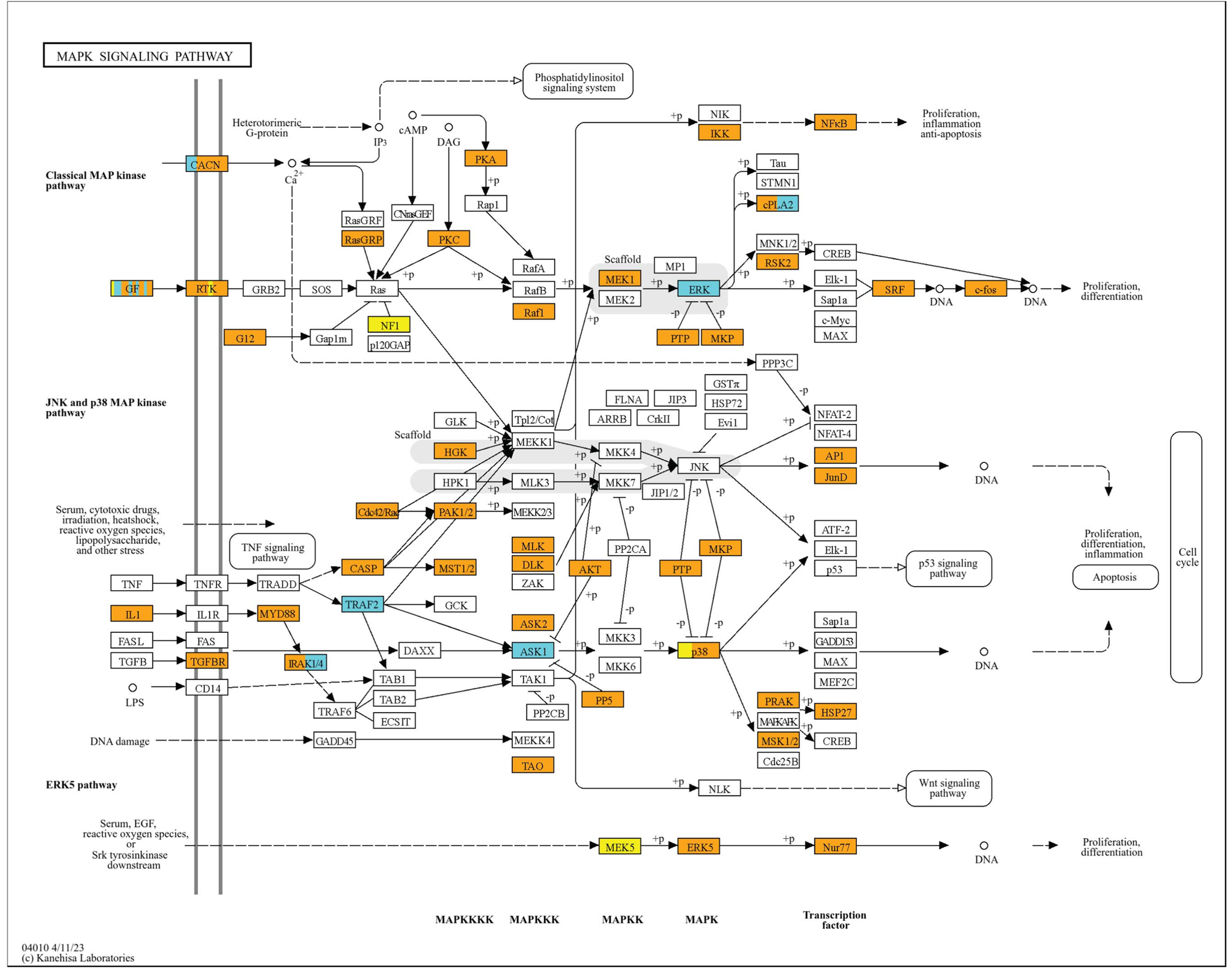

**Supplementary figure 4**: Activated THP-1 monocytes were stimulated with 150 mM ethanol, 150 mM ethanol + 100 ng/mL LPS, and 10% SAH plasma respectively. These cells were subsequently treated with 70 nM PS1145, 30 nM PH797804, or 25 µM resveratrol for 24 hours and the expression of key inflammatory and anti-inflammatory markers was analyzed.

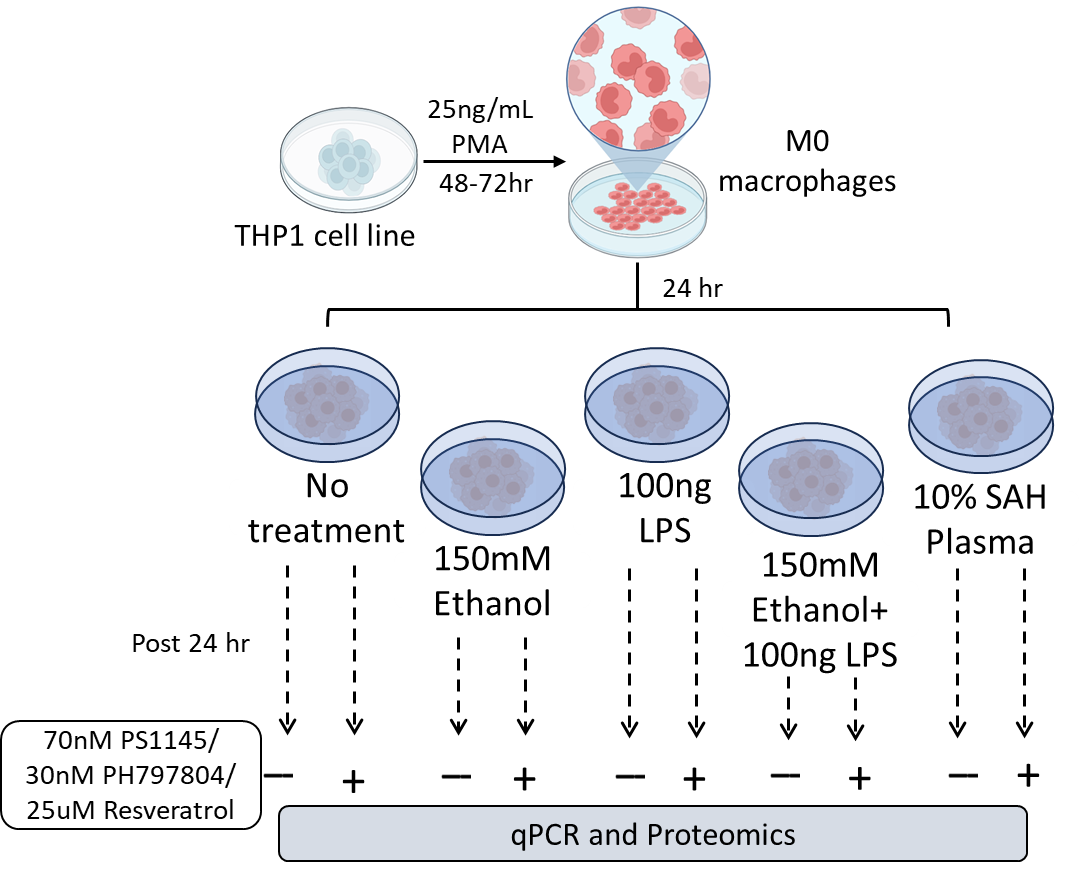

**Supplementary figure 5:** Dose standardization

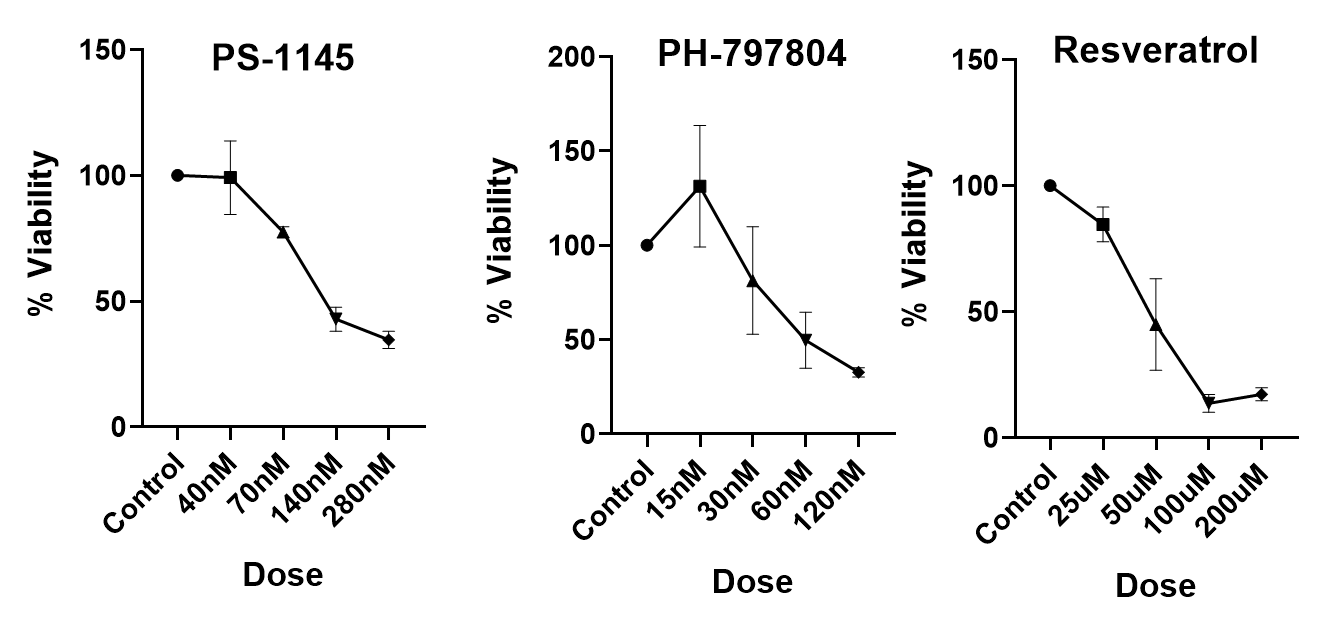
